# Myosin filaments reversibly generate large forces in cells

**DOI:** 10.1101/296400

**Authors:** James Lohner, Jean-Francois Rupprecht, Junquiang Hu, Nicola Mandriota, Mayur Saxena, James Hone, Diego Pitta de Araujo, Ozgur Sahin, Jacques Prost, Michael P. Sheetz

## Abstract

We present high resolution experiments performed on elementary contractile units in cells that challenge our current understanding of molecular motor force generation. The key features are the development of a force per motor considerably larger than forces measured in single molecule experiments, a force increase followed by relaxation controlled by a characteristic displacement rather than by a characteristic force, the observation of steps at half the actin filament period even though a large number of motors are at work in an elementary contractile unit. We propose a generic two-state model of molecular motor collections with hand-over-hand contractions and we find that these unexpected observations are spontaneously emerging features of a collective motor behavior.

---

Integrin-mediated rigidity sensing is a rich system for discovering novel features of the interplay between molecular biology and mechanics [1, 2]. In response to differential substrate rigidity, integrin based adhesions modulate a variety of downstream signals that control processes such as cell growth, death, migration [3, 4] as well as invasion [5][6] and differentiation [7].

Rigidity-sensing in early spreading cells is performed by sarcomere-like actomyosin contractile units anchored to integrins via the adaptor protein *α*-actinin [2]. These contractile units scale force production with substrate rigidity to give constant displacements. We now show on heterogenous rigidity landscapes with up to twenty-fold differences in rigidity that contractions are of the same rate (3 nm.s^*−*1^) and strain (around 60 nm) per pillar with a nearly constant overall density of active myosin heads. Over rigid pillars, the observation of a fixed strain implies the development of a large force per motor, which can be considerably larger than the forces previously measured in single molecule experiments. Here, we implemented a method to count the number of active motor heads within the contractile unit; we find that, over rigid pillars, the maximal exerted force exceeds 40 pN per myosin head. This is nearly an order of magnitude larger than measured by *in vitro* experiments (in the 1.3 to 3.7 pN range, see [1] and references herein), although it has been shown a bipolar myosin filament with only few myosin heads engaged can produce around 30 to 50 pN [8]. We interpret this result as due to the presence of tropomyosin that slows down the motion of myosin and promotes stronger myosin-actin binding forces, both effects we show are favorable to higher force generation.

By developing a new look at a generic two-state model of molecular motor collections with suitable modifications, we show that these unexpected observations are emerging features of collective motor behavior. We first show how non-muscle myosin filaments should be expected to be capable of generating very high forces at low contraction velocities. We further show that the collective behavior can, under appropriate circumstances, involve avalanche like processes which appear experimentally as steps of half the actin filament period. Our generic model provides a natural explanation for the similarity of the observed contraction and relaxation phases. It is related to the existence of a spontaneous oscillation transition in the acto-myosin system [9–14]. Here, we show that each myosin motor can resist external forces significantly larger than forces measured from single molecule experiments. We propose that the observed characteristic displacement of the pillars is due to a sharp increase in the force opposing the motion when the actin filament extremity hits the receptor tyrosine kinase AXL [15].

Previous studies have shown that the contractile units can contract in steps of half of the actin filaments repeat distance of 2.5 nm [2]. While the proposed two-state model presented here encompasses this result, it also predicts symmetric 2.5 nm steps within the relaxation phase. Here, we utilized a fast imaging camera (100 Hz) to gain the precision required to test this prediction.

Early investigations into cellular responses focused on uniform rigidity substrates. In early fibroblast spreading on fibronectin matrices, the onset and mechanisms of rigidity sensing have been well characterized [16]. After an early phase of fast, isotropic spreading that is rigidity independent, cells enter a phase of periodic protrusions and retractions of the cell edge (10-15 minutes after contact with the substrate). During this phase, sarcomere-like contractile units form in lamellipodia and pinch the substrate via nascent integrin adhesions [15, 17, 18] (Fig. 1 a,b).

**Figure 1:**
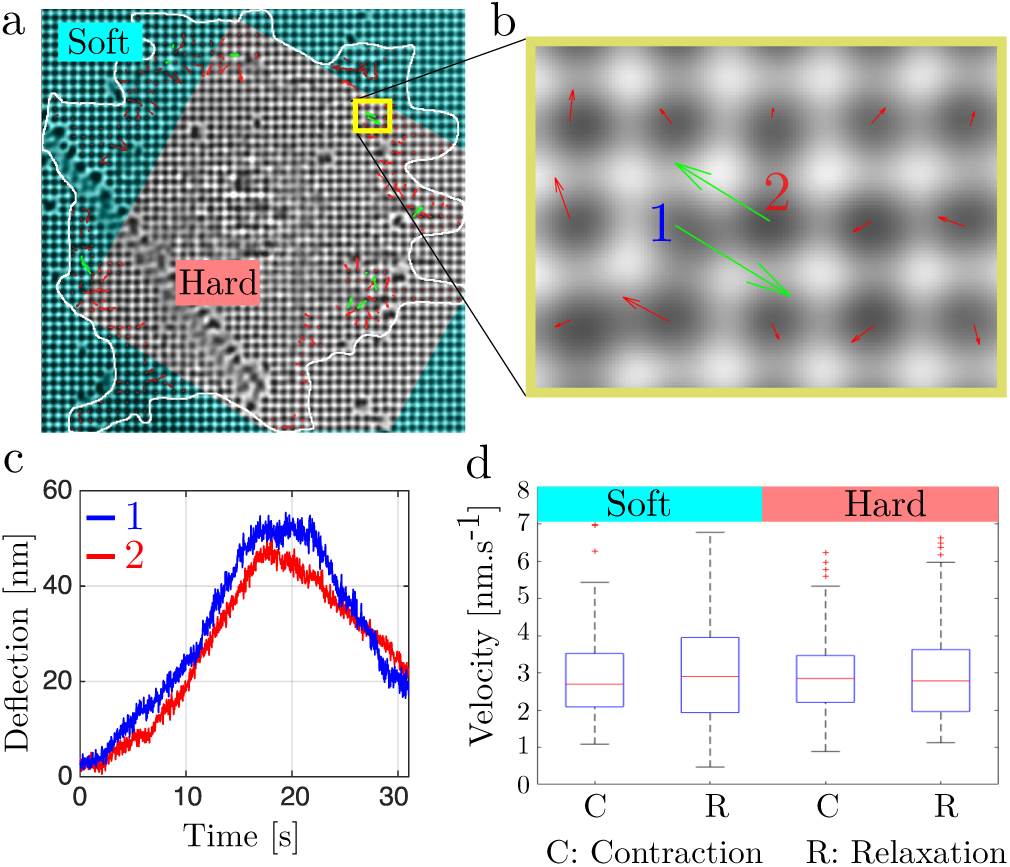
(a) Bright-field image of a cell spanning over rigidity separation lines (white line: contour of the cell) with super-imposed pillar displacement field: (red arrows) uncoordinated pillar displacements and (green arrows) displacement vectors for anti-correlated pillar pairs. The cyan-shade region marks the region with softer pillars. (b) Blow-up on (a) showing anti-correlated pillars denoted 1 and 2. (c) Pillar deflection traces of the anti-correlated pillars 1 and 2 from (b). (d) Single pillar displacement velocities measured during the contraction and relaxation phases, both for soft (*K* = 3 pN.nm^*−*1^) and hard (*K* = 60 pN.nm^*−*1^) pillars (see Method for averaging procedure).

One revealing aspect of cell rigidity sensing is that cells increase the traction forces applied to a substrate in response to increasing rigidity. Contrary to what one could have expected naively, on uniform rigidity substrates, the sarcomeric units contract to a total pillar displacement of about 120 nm regardless of bending stiffness over about a 100-fold range in rigidity [15].

In order to obtain unambiguous results regarding the number of motors per contractile unit as a function of the substrate rigidity, we created pillar-coated substrates composed of two regions with different rigidities. We focussed our attention on pillars of 500 nm diameter, since cells behaved on those pillars as if they were on a continuous surface whose elastic modulus corresponded to the beam pillar elasticity. We developed a novel method to create sub-micron PDMS micropillar arrays with distinct spatial domains of controlled rigidity, which we used to probe the rigidity sensing mechanism at the sub-cellular level.

## Experimental results

*Rigidity Responsive Tension Generation is Highly Localized and Tightly Regulated During Early Cell Spreading* In early fibroblast spreading on fibronectin matrices, rigidity sensing contractions were well characterized [2, 16, 19]. After an early rapid spreading phase, sarcomere-like contractile units formed in lamellipodia and pulled pillars toward each other for about a minute [15, 17, 20] (see Fig. 1). Unlike stress fibers, which formed after early spreading and spanned tens of microns, these sarcomere-like contractile units appeared within 10-15 minutes of contact with the substrate, and spanned *<* 4 microns. A computer program identified contractile pillar pairs using the criteria that pillar displacements of *>*20 nm were anti-parallel, *>*20s in duration and peaked at roughly the same time.

Mouse embryonic fibroblasts (MEFs) spread normally over rigid and soft areas with patterned pillar substrates (pitch: 62 *µ*m; bar width: 25 *µ*m; hole width: 37 *µ*m) with fibronectin coating (see Fig. 1a). We designed a deep UV treatment which allows for a tunable increases in the pillar bending stiffness up to twenty fold, from *K* = 3 pN.nm^*−*1^ to *K* = 60 pN.nm^*−*1^ (see Methods section). Contractile units had the same maximum pillar displacement of about 60 nm both on hard pillars (with rigidity *K* = 60 pN.nm^*−*1^) and on soft pillars with rigidity *K* = 3 pN.nm^*−*1^ (see SI Fig. S4). Further, the contraction and relaxation trajectories of single pillar occurred at nearly constant velocities of around *v* = 3 nm.s^*−*1^ both for soft and hard pillars (Fig. 1d). Thus, the contractility of these sarcomere-like units was an inherent property of the contractile units and was not influenced by substrate rigidity. Surprisingly, a twenty-fold increase in the force on the contractile units did not alter the kinetics of contraction at this slow velocity, since most myosin contractions showed a characteristic force-velocity relationship that supposedly reflected the effect of force on the release of myosin-ADP heads from actin [21–24]. However, the sarcomeric units at the leading edge of spreading cells showed no such velocity dependence on load and the unloading rate was similar to the loading rate. Thus, it appeared that a factor other than force controlled the velocity of contraction, and previous studies suggested that tropomyosin2.1 restricted the velocity of movement. To dissect the contractions in more detail, we measured pillar displacements at high temporal and spatial resolution (100 Hz). Using a previously characterized step analysis program, individual myosin stepping events were characterized in traces of pillar displacement (see SI Fig. S3e,f). As a control for over-fitting noisy data, faux displacement traces (a polynomial fit of the actual displacement trace with added pillar displacement noise from pillars outside the cells) were similarly analyzed (SI Fig. S3g,h). The analysis of faux traces gave step sizes that were fit by a gamma distribution. However, when the actual pillar displacement traces were fit, the distribution of step sizes had a prominent peak centered around 1.2 nm on top of the background noise peak (SI Fig. S3i). Somewhat surprisingly, myosin stepping size was independent of pillar stiffness (see Fig. 4), and hence resistance load, in contrast with prior in-vitro findings [25]. Further, the same 1.2 nm step size was found during both the contraction and relaxation phases on hard and soft pillars. (SI Fig. S3 l-m) Thus, a total 2.4 nm step-wise displacements (since both pillars move simultaneously [18]) were a constant feature of contraction and relaxation.

**Figure 2:**
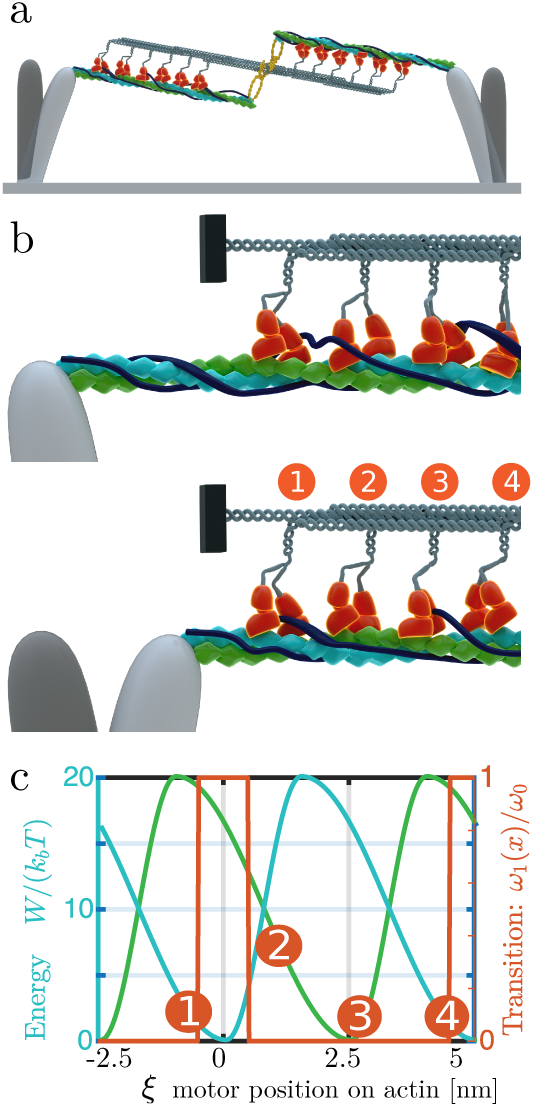
Sketch of the model. (a) Working model of contractile units, composed of doubled stranded actin filaments (green and blue helices) contracting under the effect of a myosin heads (orange). An AXL tyrosine kinase (yellow) crosslinks both the myosin thick filament to the actin strands. (b) Blow-up on the left-hand side pillar; the four represented myosin motors are not synchronized to be within the same configuration (c) Proposed hand-over-hand model: at any time, one head is bound while the other is moving towards the next binding site. The situation is described by two shifted potentials *W*_1_ (blue line) and *W*_2_(green line) describing the global interaction of the two heads with actin, which are considered to be functions of the coordinate *x* that represents the position of the actin filament with respect to the myosin tail (point of connection to the backbone); (*ω*_1_, orange curve) in simulations, we consider a transition rate that is maximal in the region of minimum configuration energy; this models the observation that myosin binding sites favor ATP hydrolysis. During a step, a myosin head initially at the rear takes the lead, e.g. the motor with configuration 1 eventually reaches configuration 3.

**Figure 3:**
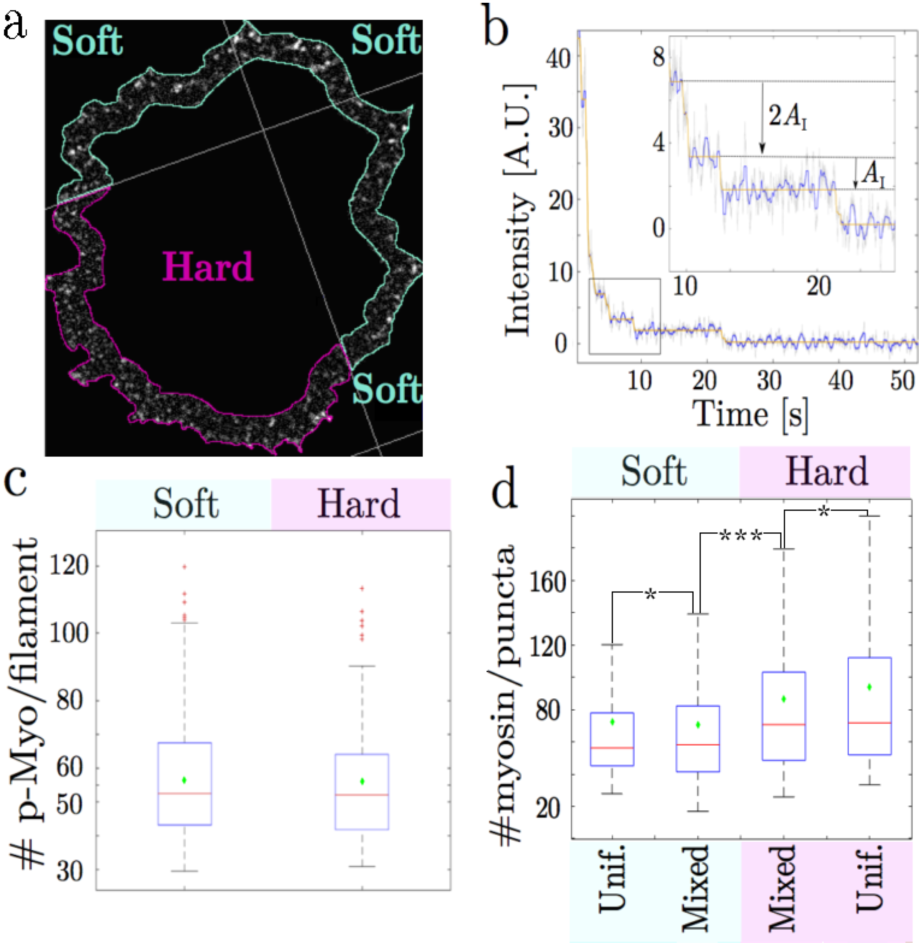
(a) pMLC staining in the lamellipodium of a cell spanning a rigidity boundary. The density of myosin filaments is uniform regardless of substrate rigidity. (b) Bleaching trace of a p-MLC punctum over 6000 20ms exposures (raw data in grey, sliding median filter in blue, step fitting in orange). (b-insert) focus on the indicated part of the curve – double and single bleaching events are highlighted. (c) Average myosin filament size in the lamellipodium for cells spread on soft pillars (8.6 kPa) and cells spread on continuous PDMS (6 Mpa): (red line) median and (green diamonds) mean values. The mean number of myosin per filament is approximatively 56 in both conditions (±2). Total number of filaments is *n >* 168 obtained form at least 10 cells for each substrate rigidities (d) Analysis of all lamellipodial p-MLC puncta shows that the proportion of myosin in multi-filament arrays is greater on more rigid surfaces, regardless of if the surface is uniform or dual rigidity. Only 1/5 of the puncta over soft surfaces have more molecules than two standard deviations above the single filament mean. Over hard surfaces 1/3 of the puncta have above two standard deviations over the single filament mean. Red lines-median, green diamonds-mean. ^*∗*^: *p >* 0.05, ^*∗∗∗*^*p*= 0.034. Soft pillar: *K* = 3 pN.nm^*−*1^; Hard pillar: *K* = 60 pN.nm^*−*1^. Continuous stiff surface here is flat PDMS, not a uniform surface of stiff pillars. The lack of a difference in puncta size between stiff pillars and flat PDMS suggests our stiff pillars are above the stiffness level at which cells can differentiate rigidity.

**Figure 4:**
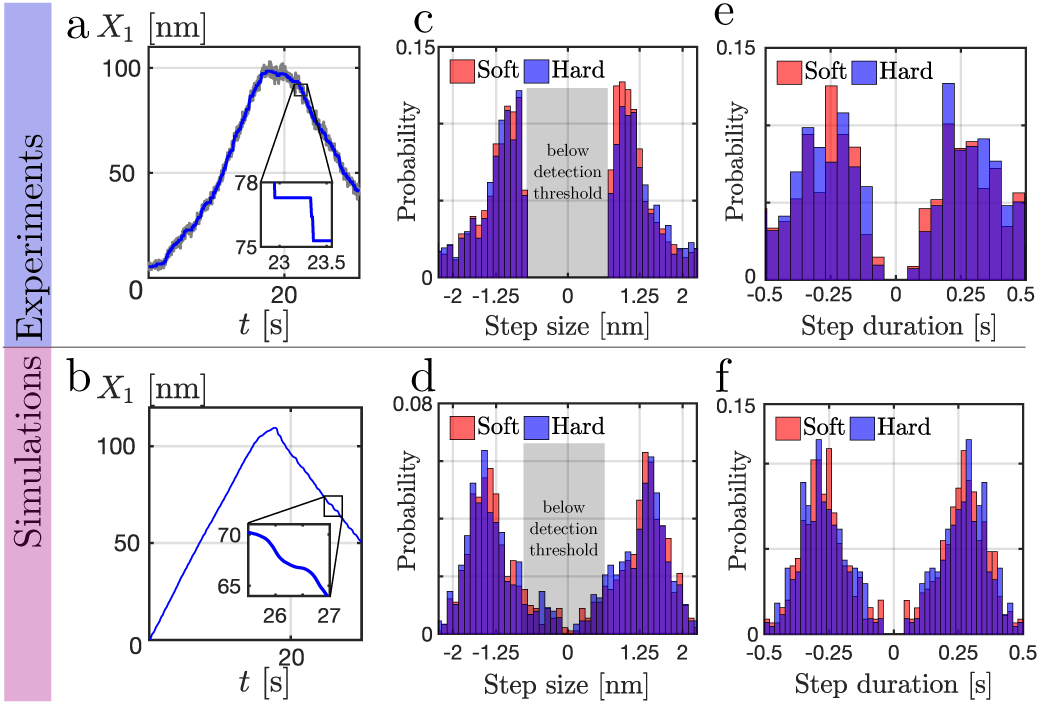
Comparison between experiments (top row) and simulations with localized transition rates at the minimum of the energy (bottom row). Left column (a, b): total deflection of the two pillars with time. Middle column (c, d): histogram of the step length for soft (red) and hard (blue) pillars. Negative values indicate steps during the relaxation phase. Right column (e, f): histogram of the duration between two steps. Soft pillar: *K* = 3 pN.nm^*−*1^; Hard pillar: *K* = 60 pN.nm^*−*1^ (see other parameter values in Tab. III).

*Myosin Filament Formation and Density is Independent of Substrate Rigidity* The invariance of speed and step length with rigidity, contraction and relaxation was possibly due in part to variations in myosin activity, since inhibition of Rho-kinase, which phosphorylates myosin light chain, abrogated integrin-based rigidity sensing [4]. To investigate the activity of myosin in the lamellipodia, we fixed cells during early spreading (20-30 m) and stained for myosin light chains doubly phosphorylated on T18/S19, which marks the highest activation state for non-muscle myosin II [26]. Image processing revealed local puncta of phospho-myosin staining in the periphery, indicative of single myosin filaments (Fig. 3a). The number of active myosin molecules per filament was determined by bleaching puncta until quantized steps in fluorescence intensity levels revealed the intensity of a single fluorophore (Fig. 3b). Using the secondary antibody labeling density, the number of active myosin heads in each bipolar myosin filament was calculated to be around 56 bisphosphorylated myosin heads on average in each bipolar filament, or 28 per half-filament (Fig. 3c; see also SI Fig. S8). This number was in perfect agreement with prior electron microscopy measurements of human platelet myosin mini-filaments [27]. This number was the same for filaments in regions on soft pillars and in regions on continuous PDMS, and was thus invariant over 3 orders of magnitude in rigidity (Fig. 3c). The fluorescence signals of filament puncta were analyzed for area and length variations in soft and rigid regions, which further indicated that single puncta were single myosin bipolar filaments with an average size of about 60 active myosin heads. (Fig. 3c) The density of filaments was also independent of the substrate rigidity at 0.3 ± 0.1 per square micron of lamellipodial area, regardless of the pillar rigidity whenever the local contractions occurred. After 1-2 hours, the level of active myosin was significantly higher over rigid areas and the cells moved toward those rigid areas as in durotaxis. However, there was no evidence of greater active myosin recruitment to lamellipodial regions over rigid pillars than soft at the early times when we observed local contractions. Nor was there evidence of a change in the balance of bulk lamellipodial myosin phosphorylation/dephosphorylation in response to substrate rigidity. Thus, the fluorescent antibody measurements of the number of active myosin heads per filament was consistent with previous electron microscopy measurements of the filament size in cytoplasm. In hard-pillar regions, however, the proportion of multi-filament puncta scaled with substrate rigidity. (Fig. 3d). Therefore, while local substrate rigidity did not change the level of myosin activation, it did cause changes in local arrangement of mini-filaments and caused multi-filament assemblies, which could have contributed to the higher forces necessary for peak displacements of hard pillars.

*Myosin Filaments Associated with Active Sarcomeric Units* To determine if there was recruitment of multiple filaments to the rigid pillar contraction units, cells on dual rigidity platforms were monitored until they reached the rigidity sensing phase of early spreading and then fixed. This enabled measurements of both pillar displacements and the number of active myosins so as to have a snap shot of the forces applied to the substrate as well as active myosin distribution at the moment of fixation. In soft pillar regions, single active filaments of average size pulled softer pillars to their maximal displacement in contractile units. In contrast, on hard pillars, multiple filaments (2-3 filaments with 100-170 active heads) were associated with contractile units at peak contraction (see SI Table II and Figs. S5-S6-S7). Thus, the high force on the rigid pillars at maximum contraction (3600 pN) was generated by 100 to 170 active myosin heads. Since the bipolar filaments were symmetrical, this force was born by half that number of active myosin heads giving a maximum force of 40 *−* 60 pN per head.

## Model

### Model: observations to be interpreted The above described results raise the following theoretical challenge

1. why is the deflection curve almost symmetrical, with nearly opposite velocities, during the contraction and relaxation phases?
2. why is the amplitude of the total displacement prescribed at a fixed value (120 nm), irrespective of the pillar elasticity?
3. why is the force exerted per motor so large compared to *in vitro* estimates?
4. why do the pillar contractions occur in steps, even though the number of motors is large (in the 100 range)?
5. why do steps correspond to half an actin period and with an essentially Gaussian distributed time interval?

*General framework* To address the challenges listed above, we adapted the two state model introduced earlier by Julicher et al [11]. This formalism provided a simplified view of the multiple step-wise chemical reactions involved in the ATP-cycle (see [10, 28]). As was known, non-muscle myosin II filaments were composed of two sets of myosin heads, which interacted with anti-parallel actin filaments, and were connected by a heavy chain domain, which bundled myosin tail domains to form a rigid backbone (equivalent to the thick filament in sarcomeres). In the following, the position of a myosin is labeled by its coordinates *x* corresponding to the contact point between this myosin and the backbone (see Fig. 2). We model its interaction with the double stranded actin filament (strands 1 and 2) through spatially varying potentials *W*_1_(*x*) and *W*_2_(*x*) which include the binding and motor conformational changes in free energies. Since each actin strand in the filament is composed of repeated G-actin monomers at *l* = 5 nm intervals along the filament direction [29], the potentials must follow the periodicity: *W_s_*(*x*) = *W_s_*(*x* + *l*), where *s* = 1, 2. In addition, as the two strands are shifted by half an actin period, the potentials must also satisfy the relation *W*_1_(*x*) = *W*_2_(*x* + *l/*2). To model the effect of ATP-binding, which provides the required energy to displace the myosin motor from one binding site to the next, we assume that a motor on the strand 1 can, independently of the other motors, undergo a transition to the other strand 2; such a transition occurs with a probability per unit time denoted *ω*_1_. We consider that the rate *ω*_1_ is maximal where *W*_1_ is minimal. Similarly, the transition rate *ω*_2_ (from strand 2 to strand 1) is maximal when *W*_2_ is minimal. The rates do not satisfy detailed balance as required for a non-equilibrium process.

*Collective assembly* Given the high elastic rigidity measured for the thick filament in muscle sarcomeres [29], we suppose that the myosin backbone is sufficiently rigid so that the distances between separate myosin tails remain fixed during the experiments. We further consider that myosin motors are distributed at regular intervals: the tail of the *j*-th motor is at the position *x_j_*(*t*) = *X*(*t*) + *q*(*j −* 1), with 1 ≤ *j ≤ N* and *N* is the total number of motors; *X*(*t*) is the position of the first motor tail. Electron microscopy studies of skeletal muscle further indicate that *q* = 14.3 nm [30]. Since *q* is not a clear multiple of *l*, the myosin and actin filaments do not seem to be commensurate with each other. We assume that the ratio *q/l* is irrational so that the sequence {*ξ_j_*} of cyclic coordinates *ξ_j_*= *x_j_* mod (*l*) covers the interval [0, *l*] with uniform density in the limit *N* = ∞ (equidistribution property).

*Dynamics* The dynamics of the tail position *X*(*t*) results from force balance

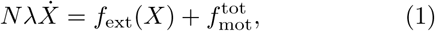

between: (i) the friction force 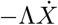, which is resisting the motion of the actin filament; (ii) the externally applied force *f*_ext_(*X*), which includes the contribution of the pillar elasticity, and (iii) the sum of the forces exerted by each motor on the myosin backbone 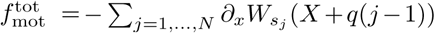 where *s* _*j*_= {1, 2} refers to the conformation of the *j*-th motor. We chose a system of coordinates such that *X* = 0 in the absence of elastic strain and such that *f*_ext_(*X*) ≈ −*KX* for small pillar deflection, where *K* is the pillar spring constant.

For a wide class of potential and transition rate functional shapes the maximum force per motor *f*_max_ is to be expected to be of the order of the maximal energy difference (i.e. 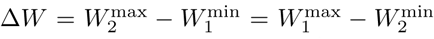) divided by the length of variation (i.e. half the period *l/*2), leading to a total force

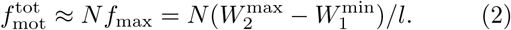

In the SI, we further derive a relation of this type for a particular model choice (see Eq. (S29)). If we now take the energy difference in Eq. (2) to be of the order of the chemical potential difference between an ATP molecule and its hydrolysis products *δµ* ≈ 20 *kT*, the force amounts to *f*_max_ ≈ 30 pN which is in the range of the forces observed here. However, as further justified in the SI, the energy difference ∆*W* could well exceed *δµ*, leading to even larger forces.

*Simulations* We calculate the dynamics of Eq. (1) by a first order discretization scheme: *X_t_*_+∆*t*_= *X_t_* + (*f*_ext_(*X_t_*)+ *f*_mot_)∆*t/*Λ, where the time increment ∆*t* satisfies *ω*∆*t* 1, to account for the stochastic dynamics of transition between strand 1 and 2 [31]. We consider asymmetric energy profiles which favour motion of the motors towards the (+) end of the filament (see Fig. 2 and SI for implementation). We choose a realistic set of parameters describing the experiments (see SI Table III). Similar to experiments, we find that the velocity of the pillar deflection curves is nearly constant in both the contraction and relaxation phases (see Fig. 4b). We further find that the force per motor is as high as found experimentally (see SI Fig. S9b).

The contraction-relaxation process appears as one period of the oscillations predicted in [10, 11] and observed in skeletal muscle myofibrils [9] and in vitro actomyosin systems [12].

In the simulations where the external force is simply the force exerted by pillars, the value of the maximum displacement depends linearly on the pillar spring constant: what determines the cross-over between contraction and relaxation is a force, not a displacement. The latter statement clearly disagrees with experimental observations. However, the force acting on filaments may not be due to pillars only. In particular, previous experiments have shown the crucial role of the receptor tyrosine kinase AXL in regulating the maximal pillar deflection [15]. It is very likely that beyond a certain displacement, steric hindrance restricts the motion of the filaments. Here, we consider that AXL contributes to a non-linear elastic term, changing the external force from *f*_ext_(*X*) = *−KX* for low deflection strains *X < l*_AXL_ to *f*_ext_(*X*) = *−K*_AXL_*X* for larger deflection strains *X > l*_AXL_, where *K*_AXL_ *≫K*. We set *l*_AXL_ = 120 nm. With this improvement, our simulations indicate a maximal summed displacement *X*_*max*_ ≈ 120 nm independently of the pillar elasticity *K*, again in good agreement with experiments (see Fig. S9a).

Moreover, we observe the presence of steps with *l/*2 actin period in the simulated trajectories (see Fig. 4), again in good agreement with the experimentally observed 2.5 nm steps in the total pillar deflections. In the SI Movie 1, we show that the step dynamics is reminiscent of an avalanche process. The motion of the motors alternate between fast and slow regimes of contractions: during the slow regime, motors in the low energy state (i.e. at a binding site) undergo a transition to the higher energy state (i.e. via consumption of an ATP); such progressive increase in the energy of the assembly will be dissipated during the following fast regime. The step length and waiting time statistics are similar to the ones observed in experiments (see Fig. 4).

Thus simulations agree nicely with all observed original features apart from one exception: the theoretical results indicate that there will be a periodic behavior of contraction-relaxation phases, whereas in the experiment there is only a single contraction-relaxation event. This difference is due to the fact that when the applied force vanishes, the contractile unit disassembles. We will address this aspect in the discussion section of the manuscript.

The question is now to get a proper analytic understanding of the results.

*Analytical Framework* We consider an analytical treatment which corresponds to the limit of a large number of motors *N* ≫ 1. In this limit, the motor population *N*_1_ in state 1 can be expressed in terms of a motor density *P* _1_ as 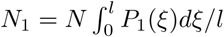, where *ξ* = *x* mod *l*; similarly, we define *P*_2_ as the motor density in state 2. The equidistribution of the sequence of cyclic coordinates {*ξ_j_*} leads to the relation: *P*_1_(*ξ*) + *P*_2_(*ξ*) = 1*/l*. The Fokker-Planck equation relating the evolution of the occupancy probability distribution *P*_1_ then reads:

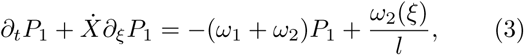

where the velocity of the myosin filament 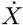 is to determined through the force balance equation Eq. (1).

The question is now whether or not one can observe the steps in the large number of motors limit of Eq. (3). For that aim, we seek an analytical solution of Eqs. (1) and (3) by expressing the *l*-periodic function *P*_1_ as

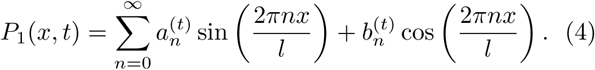

In the general case where *ω*(*x*) = *ω*_1_(*x*) + *ω*_2_(*x*) is not uniform, the term *ω*(*x*)*P*_1_ in Eq. (3) results in a product of different Fourier components, which leads to an infinite system of equations on the coefficients (*a_n_, b_n_*). To make progress, we simplify the problem by truncating the energy and transition rate profiles to their first mode: *W*_1_(*x*) = *W* (1 − cos(2*πx/l*))*/*2, *ω*_1_(*x*) = *ω/*2 (1 + cos(2*πx/l*), *W*_2_(*x*) = *W* (1 + cos(2*πx/l*))*/*2 and *ω*_2_(*x*) = *ω/*2 (1 − cos(2*πx/l*) such that the total transition rate *ω*(*x*) is constant [32, 33].

Under these assumptions, we obtain a numerical solution to Eq. (3) for the evolution of the probability distribution *P*_1_(*x, t*). The resulting numerical trajectory *X*(*t*) displays steps with *l/*2 period, which shows that the stepping behavior is also preserved in the limit of a large number of motors. The SI Movie 2 provides an intuitive interpretation for the steps. There, the initial motor repartition (*NP*_1_) is mainly in the negative slope of the energy, corresponding to a strong positive motor force which further increase the velocity of the assembly. As the velocity increases, the density profile *P*_1_ is advected in the positive *x* direction; this progressively increases the fraction of motors that are in the positive slope of the energy, further contributing to a net decrease of the motor force 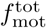) hence to a decrease in motor assembly velocity. Such decrease in the motor assembly velocity favors the progressive shift of the motor density from the low-energy regions towards higher energy regions – which provides the energy required for a subsequent motor step. Such a delayed negative feedback-loop between the motor assembly velocity and force naturally leads to oscillations [34].

We now prove that the distance covered during an oscillation of the velocity is exactly *l/*2. For simplicity, we focus on the case of a vanishing restoring force *f*_ext_ = 0. As represented in Fig. 5a, the time evolution *X*(*t*) exhibits steps, but only at short times *ωt <* 3. At longer times, the steps disappear as the velocity becomes nearly constant. As further justified in SI, we find that the dynamics of the variable *y* = ln(*a*_1_*l*) is given by the equation:

**Figure 5:**
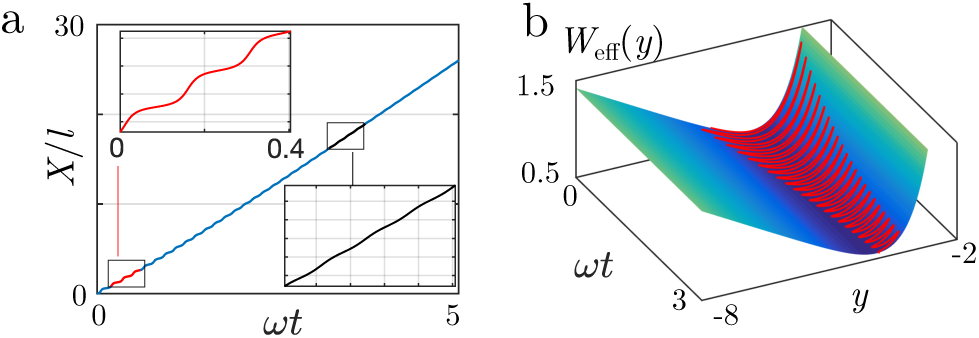
Analytical description for the 2.5nm step behavior. (a) Displacement as a function of time, in units of *l*, in the absence of an external force (*ρ* = 0; *µ* = 10^3^). The stepping behavior with half-actin periodicity is visible at short times (*ωt >* 1, red frame) but vanishes at longer time (*ωt >* 3, black frame). (b) The step behavior can be mapped into the swing motion of particle in an effective potential *W*_eff_: (solid red line) trajectory of Fig. (a) in the space (*ωt, y, W*_eff_ (*y*)). The amplitude of the oscillations in *y* weakly dampens with time; the step behavior are visible during a long-lived transient.

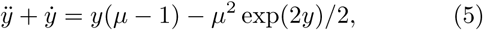

where *µ* = 2*π*^2^*W/*(*l*^2^*ωη*) is a dimensionless quantity referred to here as the *activity* parameter. Mapping Eq. (5) to the dynamics of a massive particle in an effective potential 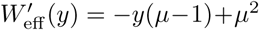, one expects trajectories *y*(*t*) to exhibit damped oscillations (see Fig. 5b). For large *µ*, we find that the typical number of observable oscillations before damping is of order *µ*, so it can be quite large. During this oscillatory transient, one can neglect the damping term *ẏ*, in which case the quantity *ẏ*˙^2^*/*2 + *W*_eff_ (*y*) is a constant corresponding to the conservation of energy in this massive particle problem; from this, we deduce the period of the oscillations, as well as the distance covered *δX* during this period. In the limit *µ* ≫ 1, we find that 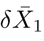 = *l*/2 (see detailed proof in the SI).

Steps correspond to a transient behavior due to the presence of the friction term in Eq. (5). Yet the transient stepping behavior can involve a large number of events in the large *µ* limit, which corresponds to the case of low transition rates and the velocities, hence to the experimentally relevant regime observed here. Indeed, the presence of tropomyosin decreases the transition rates *ω* by several orders of magnitude compared to motility assays, leading to *µ* values of order 10^3^ (see SI, as well as [32]). Interestingly, in motility assays steps are not observed for numbers of motors exceeding a few [35]. We find that the relevant *µ* values are in that case much smaller; this corresponds to the regime where damping is sizeable and such that a steady state is reached before steps can be observed.

One important conclusion is that the relaxation phase also starts as a transient state because of the initial sharp sign change of the averaged velocity. Thus, experiments should exhibit steps in the relaxation phase as well as in the contraction phase.

*Model predictions validated: step behavior during the relaxation phase* Since the theoretical analysis predicts the existence of steps in the relaxation phase, we decided to analyze the experimental relaxation curve *X*(*t*) with the same step finding algorithm as we used for the contraction. As predicted by our analysis, the relaxation steps are practically identical to those observed during contraction (see Fig. 4). They are characterized by the same average half period length and similar statistical distributions. We take this observation as a confirmation of the validity of the theoretical analysis.

## Discussion

In this work, we determine from single cells spanning regions of pillar arrays of different stiffnesses that:

- contractile units forming in the cell periphery undergo a contraction phase followed by a nearly symmetric relaxation,
- forces per motor can exceed by orders of magnitude the range of forces observed in single molecule experiments,
- the turning point between the two phases is set by a displacement linked to the presence of AXL,
- the contraction phase involves displacement steps of half the actin period, the waiting time between steps is Gaussian distributed.

Our analysis of the two-state model of molecular motor collections allows us to understand all observed features as emerging features due to the large number of motors. These results are obtained under conditions appropriate to the experiment, in particular taking into account the role of the tyrosine kinase AXL and the decreased rate of motion due to tropomyosin. Our model encompasses the early prediction of oscillatory behavior of motor collections [11] but contains new and unexpected features specific to the current situation, such as the appearance of steps as a transient behavior and the switch between contraction and relaxation determined by a length rather than a force. As predicted by our theoretical model, steps are observed experimentally within the relaxation phase with statistics very similar to those of the contraction phase.

The only sizeable difference between experiments and theory is the fact that, experimentally, contraction and relaxation occur only once whereas theory predicts a periodic set of successive events. We understand this difference as a result of the fact that the contractile unit spontaneously disassembles under a no load condition. Disassembly has to be fast enough to be completed before the force grows again. In turn this implies that many attempts at forming the contractile unit fail before one event is successful. We plan to analyze the waiting phases during which no major events are observed on the pillars: one could see the signature of attempts at creating contractile units, for instance by observing steps in the fluctuations. Eventually one can speculate on the physiological relevance of the contraction-relaxation events of the contractile units. The simplest hypothesis could be that these experiments correspond to attempts at constructing larger objects such as stress fibers. Even more interesting is the possibility of rigidity sensing: since the displacement is prescribed, the force is not, and the maximum developed tension depends on the substrate stiffness. Force dependent signaling could easily be used by cells to adapt their physiology to the surrounding [1].

## Conclusion

An unexpected result of our study is that cells pull pillars of twenty-fold different rigidities with the same velocity and over the same spatial extent, generating forces in excess of about 40 pN per myosin head on the stiffest pillars. Although these forces are much larger than the forces measured per myosin head in vitro, we present here theoretical analyses of bipolar myosin filament arrays that show that such forces are possible under low velocity conditions.

## Methods

*Cell Culture* MEFs were cultured in DMEM (Gibco) supplemented with 10% (vol/vol) FBS (Atlanta Biologicals), 2mM L-glutamine (Gibco), and 100 IU/mg penicillin-streptomycin (Sigma) at 37 C and 5% CO2. PDMS pillars substrates were coated with human plasma fibronectin (10 *µ*g/mL, Roche) and incubated at 37° C for 1 hour, then rinsed 4X with PBS to remove excess free fibronectin. PBS was replaced with Ringer buffer before plating the cells. Cells were trypsanized and suspended in Ringer buffer at 37° C for 30 minutes prior to plating on fibronectin coated pillars.

*Deep UV Treatment Allows for Tunable Increases in Pillar Bending Stiffness up to Twenty Fold* Exposure of PDMS to UV/Ozone crosslinked-Si02-groups into-Si0X-creating a 80 nm thick silica-like layer below surface of the PDMS [36–41]. This change was irreversible, and the effective bending stiffness of UV/Ozone treated pillars increased with longer exposure times as measured by Atomic Force Microscope (AFM) through repeated bending measurements (see SI Figs. S1-S2). Pillar arrays were patterned with sub-cellular regions of soft pillars and stiff pillars by placing nickel Transmission Electron Microscopy (TEM) grids on top of the pillar surface prior to UV/Ozone treatment. A fluorescent dye that was broken down by UV light (coumarin 343) was added to the PDMS to mark regions shielded from exposure (SI Fig. S1a, note blue regions were not exposed).

Although pillar bending stiffness was often calculated by the simple Euler-Bernouli beams [42, 43], the stiffness of UV treated pillars had to be measured. Using the AFM in contact mode, the tip was scanned back and forth horizontally, repeatedly bending and unbending the pillar (SI Figs. S1b-S2) while the torsional motion of a calibrated cantilever was recorded. The bending and un-bending readouts were linear and reproducible, demonstrating that pillars were Hookean springs for the displacement ranges tested (up to 100 nm). Bending stiffness increased with exposure times, reaching 20-fold after a two-hour exposure with no sign of saturation (see SI Fig. S1c). For these experiments, a two hour irradiation was used. Analysis of pillars in a line across a rigid-soft boundary showed that the rigidity transition occurred over 3 pillars (2 microns). Thus, pillars chosen for analysis were always above 2 microns from the transition boundaries where the stiffness was reproducibly measured.

*Pillar velocities estimations* The pillar deflection velocities presented in Fig. 1d are averaged over a time window [*t*_start_ + 3 s*, t*_end_ − 3 s] where *t*_start_ and *t*_end_ refer to the start time and the end time of the contraction (resp. relaxation) phase.

## Supplementary Information

### Appendix A: Experimental protocol

#### 1. Deep UV Treatment Allows for Tunable Increases in Pillar Bending Stiffness up to Twenty Fold

Exposure of PDMS to UV/Ozone crosslinks-Si02-groups into -Si0X-creating a 80 nm thick silica-like layer below surface of the PDMS. A schematic of the mask and UV/Ozone treatment can be found in Fig. S1a, with the inset image showing an example of a cell spread across multiple regions of soft and stiff pillars. Blue fluorescence marks the soft regions.

Since PDMS pillars were commonly used as a force sensing cell culture platform, pillar bending stiffness was characterized with particular regard to the assumption that they bend as simple Euler-Bernouli beams. The PDMS transition to silica was not uniform through the entire volume of a pillar; therefore, we couldnt simply plug the adjusted Youngs modulus into a bending stiffness formula to determine the spring constant of treated pillars. Instead, the bending stiffness was measured directly by an AFM based method. Briefly, the pillar tops were imaged in tapping mode in order to localize the pillar positions within the scanning area of the AFM tip. Then the AFM was switched into contact mode and the tip was brought into contact with the center of one of the visualized pillars. With the tip in constant contact with the pillar top, the tip was scanned back and forth horizontally, repeatedly bending and unbending the pillar (Fig. S1b) while the torsional motion of the cantilever was recorded. Since the torsional stiffness of the cantilever was much higher than the bending stiffness of the pillars, displacement of the pillar top was equivalent to the horizontal translation of the sample and the torsional forces the cantilever gave a direct measure of pillar bending stiffness.

The trace (bending) and retrace (unbending) readouts of torsional motion of the cantilever were linear and overlapping over repeated displacements, demonstrating that the AFM tip did not slip once brought into contact with the pillar top, and that pillars displaced horizontally in the plane of the pillar tops act as Hookean springs for the displacement ranges tested (up to 100 nm) with the slope of the torsional displacement proportional to the pillar bending stiffness. In Fig. S1c, we show the increase in bending stiffness for different pillar heights and exposure times, with the greatest increase topping 20-fold after a two hour exposure. There was no saturation in stiffness increase with up to two hours of treatment; and hence, greater increases were possible.

Pillars in a line across the rigid-soft boundary were analyzed to determine how sharp the transition was from rigid to soft. In narrow regions, pillar stiffness was measured for transitions from hard to soft and back to hard (Fig. S1d). With pillars spaced 1 micron center to center, the rigidity transition occurred over 3 pillars (2 *µ*m). Additionally, on surfaces where rigid areas were 25-35 *µ*m wide, transition pillars constituted less than 10% of the total.

#### 2. Pillar Displacement Tracking Microscopy

Time lapse imaging of pillars was performed with bright-field microscopy using an Orca-flash 2.8 camera (Hamamatsu) attached to an inverted microscope (Olympus IX-81) maintained at 37° C with a temperature isolation chamber running MicroManager software (UCSF). Images were recorded at 1 Hz using a 100x objective (1.4 NA oil immersion, Olympus). The centroid of each pillar was calculated using the NanoTracking [18] plug-in for ImageJ software (National Institutes of Health). The time-series positions of all pillars in contact with the cell were fed into a MatLab program that subtracted off stage drift by using untouched pillars as fiducial markers, removed noise from the displacement traces using a Butterworth filter, and identified contractile units based on thresholds of separation (nearest or next-nearest pillars), directionality (less than 90 degrees away from anti-parallel displacement vectors), displacement (both pillars exceed 30 nm displacement), and duration (contraction exceeds 15 s). The program also generated pillar displacement videos that highlight contractile units in real time (S. Movie 3).

#### 3. High Speed Pillar Tracking

High speed (100 Hz) images were acquired on the same equipment as the pillar displacement tracking with the following changes. The temperature isolation chamber was lowered to 23° C in order to slow myosin motor ATPase activity and ratcheting action to better resolve individual steps. The condenser shutter was removed to eliminate noise, and a 600 nm high pass filter was added to the light path to increase the light intensity without incurring photo-damage to the cells. Acquisition was taken in continuous 10 ms exposures. Camera acquisition was limited to a 400×600 pixel ROI so that the data from the camera would transfer fully during each acquisition frame and not cause a slow down in acquisition, giving us the highest time resolution possible. Displacement traces were smoothed using a sliding 15-point median filter. Previous work showed that steps occur every 0.25 s (25 frames) at 37° C, therefore this filter will not blur consecutive steps, especially for contractile units at lower temperature. Smoothed traces were fit using the step finding algorithm L1-PWC [44]. This algorithm generated both distinct steps, where two steady displacement values were separated by an abrupt change (more than 1nm), and ramps, where many consecutive short lived plateaus in displacement where separated by very small distances (less than 0.5 nm). The ramps likely represented over-fitting of the data. To differentiate between real steps, and over-fitting we generated an artificial data set for each pillar displacement analyzed. To accomplish this we fitted a polynomial to each pillar trace, and then overlaid onto the polynomial fit the noise in our system (the measured displacement of a pillar that was not in contact with the cell and thus experienced no applied force). The L1-PWC algorithm fit these artificial data sets mostly with ramps. In order to partially offset the over-fitting, steps of less than 0.3 nm were eliminated by combining the plateaus on either side and averaging the displacement of all time points contained in the new plateau. After this correction was applied to both the measured traces and the generated artificial traces both sets of data (real and artificial) showed a background distribution of steps that could be well fit by a gamma distribution. However, for the measured data there was also a clear contribution of defined steps in these distributions. Therefore, we cannot identify every stepping event in a pillar displacement, but we can identify and characterize the ensemble of myosin-driven stepping events.

#### 4. Immunostaining

Cells were fixed for 12 minutes with 4% formaldehyde solution (Sigma, diluted from 37% in PBS). They were then permiabalized for 4 minutes in 0.2% Triton X-100 (V/V, Sigma), and blocked in 1% BSA (W/V, Sigma) at room temperature for one hour. Primary antibody (Phospho-Myosin Light Chain 2 (Thr18/Ser19), Cell Signaling Technology) was diluted 1:100 in 1% BSA and incubated on samples overnight at 4° C. Secondary antibody (Anti-Rabbit IgG-Alexa 647, Invitrogen) was diluted 1:500 in 1% BSA and incubated on samples 1 hour at room temperature.

#### 5. Phospho-Myosin Molecule Counting

Immunostained samples were visualized on an inverted microscope (Olympus IX-81) with a 60x objective (1.45 NA) oil immersion, Olympus, an EMCCD camera (Cascade II: 512, Photometrix), and an additional 2X magnification (Spot Imaging Solutions DE20TMT). A bleaching time course is obtained by focusing in the plane of the pillar tops (the ventral surface of the cell) and imaging 6000 consecutive 20 ms exposures with epi-illumination. In order to isolate individual bipolar filaments from out of focus background illumination, we applied a 667 nm radius rolling ball background subtraction to the image sequence. It is necessary to analyze only well separated filaments, both so that the rolling ball does not artificially dim the signal due to nearby filaments, and so that we are only counting single filaments when we look at number of molecules per filament. Since there are fewer than 0.5 filaments on average in the area of this rolling ball filter, there are enough isolated filaments to acquire multiple bleaching curves per cell. Bleaching curves are obtained by taking the average intensity of a 5×5 pixel square surrounding an isolated filament at each time point. 5 pixels corresponds to 667 nm, which is larger than a single NMM2 filament. This size allows for some stage drift during acquisition without losing any signal. These curves are then smoothed with a 50-point sliding median filter and fitted with the L1-PWC step finding algorithm. Noise levels are still high compared with step sizes after smoothing, so to ensure accurate measures of single fluorphore signal level we only count differences between steps that last for at least 300 frames. For instance, in the curve in Fig. 3b we would only count two steps, because the first step is not preceded by a steady state of more than 300 frames. Additionally, steps are only counted if they are separated by over 2X the standard deviation of the noise. The noise is defined as signal after the curve reaches its final steady state (no signal). The step size extracted from these bleaching curves (multiplied by 25 because it is the average of a 5×5 pixel square) tells us the signal due to a single alexa fluorophore. We can then determine the number of myosin molecules in a mini-filament by dividing the fluorescence signal of the filament by the signal due to a single fluorophore and the average number of fluorophores conjugated to a secondary antibody (Fig. S8 g).

#### 6. Myosin Filament Size Determination

After rolling ball background subtraction was performed on immunostained images, the lamellipodium was isolated by taking the outermost 3 *µ*m of the leading edge (Fig. 3a). In order to limit the image-to-image variability, we used the ImageJ particle analyzer to threshold all images at the same value. The intensity of all puncta between 24 square pixels (the minimum area for a bipolar filament) and 48 square pixels (representing the largest area a single filament could occupy) were recorded (Fig. S8 k). These intensity numbers were then adjusted up to compensate for any signal lost due to thresholding (weak signal at the edge pixels of a punctum is cut off due to thresholding). To adjust up to the full intensity, the maximum and threshold cut off for each punctum were fitted to the point spread function of the imaging setup (Fig. S8 j) to find the z-score of the threshold cutoff. Any area not under the point spread function at that z-score was then added to the total of that punctum to get the true intensity value. After isolating single filaments, molecule numbers were determined by dividing the adjusted punctum signal by the signal intensity for a single fluorophore and the average number of fluorphores per secondary antibody (Fig. S8 g).

## Appendix B: Theoretical appendix

### 1. Model discussion

#### a. The two-state model is generic

Here, we discuss the applicability of our two-state model; we show that it is a generic framework to understand the collective behavior of motors, independently of the detailed mechanism of motion of each motors. The main feature of our model lies in the *l/*2 = 2.5 nm translational symmetry between the two states, such that *W*_2_(*ξ*) = *W*_1_(*ξ* + *l/*2) and *ω*_2_(*ξ*) = *ω*_1_(*ξ* + *l/*2). This symmetry results from the spatial shift of 2.5 nm between the two actin protofilaments. In the following, we detail two particular models for which the comparison to a two-state model is particularly straightforward.

In the main text, we introduced the possibility that myosin heads move according to a hand-over-hand mechanism, by which the two head are perfectly synchronized (see Fig. 2). As opposed to several *in vitro* studies that have shown that the two heads of a single myosin II motor work in an uncoordinated fashion, we propose a fully coordinated mechanism analogous to the hand-over-hand model for kinesins: at any time, one head is bound while the other is drawn towards the next binding site. In this framework, we model the interaction between a myosin II bi-motor and the actin filament through a pair of spatially varying potentials *W*_1_(*x*) and *W*_2_(*x*) (see Fig. 2). The powerstroke corresponds to the conformational change by which the rear-ward head progressively takes the lead as it reaches the next binding site (see Fig. 2b). In this model, the symmetry between the two heads translates into the relation: *W*_1_(*x*) = *W*_2_(*x* + *l/*2). Another possibility, which corresponds to limit of perfectly desynchronized heads, is the *strand-switching* mechanism: each head could switch to the nearest binding site on the other strand of the filament, independently of the other one. Both models satisfy the required 2.5nm translation symmetry.

#### b. Energy barriers can exceed the energy of a single ATP hydrolysis

Following [10], it can be shown that the condition of detailed balance of the chemical reaction leads to the following expression of the transition rate from state 1 to state 2:

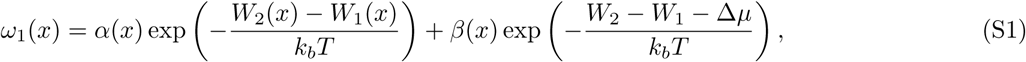

where *α*(*x*) and *β*(*x*) define the coupling between the reaction kinetics and motor conformation (parameterized by its tail position *x*); *δµ* = *µ*_ATP_ *− µ*_ADP_ *− µ*_P_ *≈* 25 *k_b_T* represents the ATP consumption energy [45]. Since

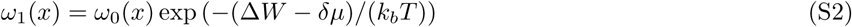

where *ω*_0_ can be expressed in terms of *α* and *β*. Following [46], we consider that the frequency *ω*_0_ corresponds to a constant given by Eyring’s theory of chemical reaction, whereby *ω*_0_ is interpreted as a molecular vibration mode: *ω*_0_ ~ *h/k_b_T* ≈ 10^12^ Hz [47] (see also [29]).

The latter relation expresses the relation between step rate *ω* and the energy barrier (∆*W* = *W*_2_ − *W*_1_) that needs to be crossed in order to perform a step. Given the experimental observation of a slow contraction of the motor assembly, we expect the typical range of the frequency *ω* to be in the range 1 Hz range. This leads us to the conclusion that the energy barrier to perform a step is in the range:

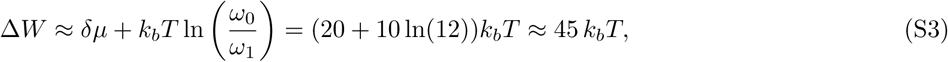

which appears to be approximatively twice larger than the value of the ATP hydrolysis *δµ*. The second hand term in Eq. (S3) suggest that thermal fluctuations can play a critical role.

The low velocity of the motor assembly is consistent with low values of the transition rate *ω*, which, in turn, is consistent with the existence of a large amplitude energy profile ∆*W > δµ*. Due to the large value of the energy gap that needs to be overcome in order to generate a motor step, we expect that a large number of ATP-consumption events correspond to unsuccessful attempts in generating a motor step. Successful motor steps could correspond to a single ATP-consumption events (in agreement with [48]) that are assisted by thermal fluctuations.

Within this energy landscape, the motors can generate larger force than usually expected. This unusual set of parameter values are likely due to the specific experimental conditions and, in particular, due to the presence of tropomyosin [20].

### 2. Numerical implementation

*Shape of the potential* We consider the class of asymmetric potential in the form:

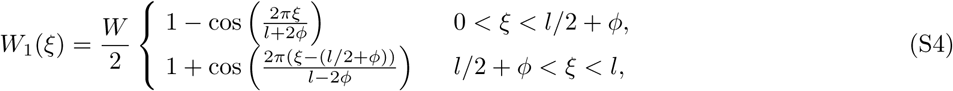

where *ϕ <* 0 corresponds to a profile with a higher slope in the + direction (see Fig. 2). The transition rate from state 1 to state 2 is taken to be localized at the minimum of the energy: *ω*_1_ = *ω*_0_ for 0 *< ξ < e* and *l − e < ξ < l*. The transition rate from state 2 to 1 then reads: *ω*_2_ = *ω*_0_ for *l/*2 − *e < ξ < l/*2 + *e*. In the simulations, we considered a potential asymmetry parameter *ϕ* = 0.30 nm, which corresponds to the value used for the Movie 1. As shown in Fig. S9b, the energy profile of Eq. (S4) can lead to the generation of large forces that can reach 60 pN.

*Dynamics* The dynamics of Eq. (1) by a first order discretization scheme: 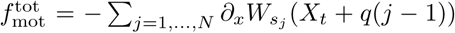 where *ω*_1_Δ*t* ≪ 1 and 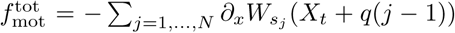 where *S*_*j*_ = {1, 2} refers to the conformation of the *j*-th motor. We consider that the external force reads *f*_ext_(*X*_*t*_) = − *KX*_*t*_ for low deflection strains *X*_*t*_ < *l*_AXL_ and that *f*_ext_(*X_t_*) = −*K*_AXL_*X_t_* for larger deflection strains *X_t_ > l*_AXL_, where *K*_AXL_ ≫ *K*. We set *l*_AXL_ = 120 nm and *K*_AXL_ = 300 pN.nm^*−1*^.

*Parameter discussion* Compared to the default set of parameters defined in Table III, our simulations indicate the following trends:

1. decreasing the transition rate *ω* leads to more visible steps but reduces the maximal force generated per motor.
2. increasing the friction coefficient *η* leads to longer contraction and relaxation phases.
3. increasing the asymmetry parameter *ϕ* leads to shorter relaxation phases, which are interrupted by a new phase of contraction before reaching *X* = 0. The origin of this behavior lies in the large speed oscillations (which correspond to the observed step behavior) which are then sufficiently large to change the sign of the velocity, thus prompting a new contraction phase. For very large values of the potential asymmetry *ϕ* (*>* 1 nm), the trajectory typically remains fixed around *X ≈ l_AXL_*: the motor assembly is unable to relax the accumulated stress.

## 3. Analytical proofs

### System of equations

We consider an analytical treatment which corresponds to the limit of a large number of motors *N ≫* 1. In this limit, the motor population *N*_1_ in state 1 can be expressed in terms of a motor density *P*_1_ as 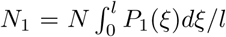, where *ξ* = *x* mod *l*; similarly, we define *P*_2_ as the motor density in state 2. The Fokker-Planck equation relating the evolution of the occupancy probability distribution *P*_1_ then reads:

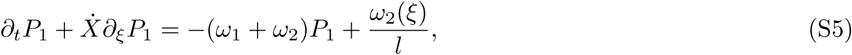

where the velocity of the myosin filament 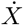 is to be determined through the force balance equation Eq. (1). In the large number of motor limit, the total force exerted by the motor assembly can be expressed as an integral over the probability distributions:

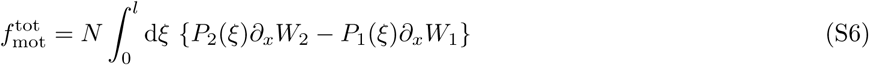

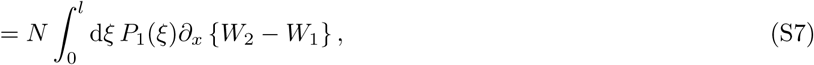

where we made use of the condition *P*_2_(*ξ*) = 1*/l − P*_1_(*ξ*).

We seek an analytical solution of Eq. (S5) and Eq. (S6) by expressing the *l*-periodic function *P*_1_ as

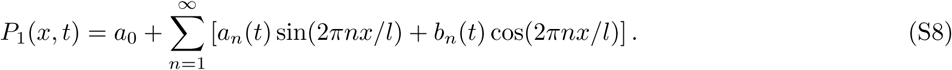

In the general case where *ω*(*x*) = *ω*_1_(*x*) + *ω*_2_(*x*) is not uniform, the term *ω*(*x*)*P*_1_ in Eq. (3) results in a product of different Fourier components, which leads to an infinite system of equations on the Fourier coefficients. Following [32], we simplify the problem by truncating the energy profiles and transition rates to their first mode: *W*_1_(*x*) = *W* (1 − cos(2*πx/l*))*/*2, *ω*_1_(*x*) = *ω/*2 (1 + cos(2*πx/l*), *W*_2_(*x*) = *W* (1 + cos(2*πx/l*))*/*2 and *ω*_2_(*x*) = *ω/*2 (1 − cos(2*πx/l*) such that the total transition rate *ω* is constant [33]. Under these assumptions, the motor forces from Eq. (S6) can be expressed in terms of the first Fourier coefficient as: 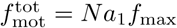, where *f*_max_ = *πW/l*.

Furthermore, inserting the Fourier decomposition into Eq. (S5), we find (i) that the first mode *a*_0_ is decoupled from the other modes and that it relaxes exponentially fast to its steady state value 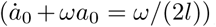, and (ii) that *a*_1_(*t*) and *b*_1_(*t*) evolve independently from the other modes *a*_0_, *a_n_* and *b_n_* for *n >* 1, with:

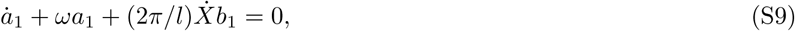

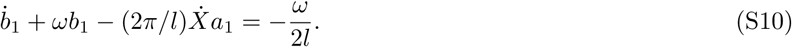

The dynamics of *X* is given by the force balance equation:

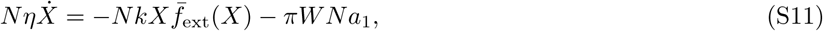

where we define *k* = *K/N*. In deriving Eq. (S11), we used that

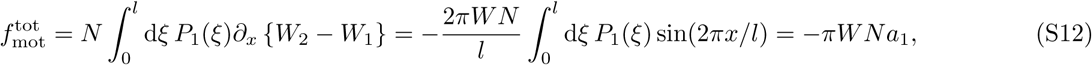

due to the orthogonality relation: 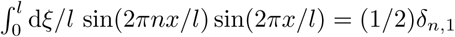. Combining Eqs. S9-S10 and Eq. (S11) leads to the dynamics:

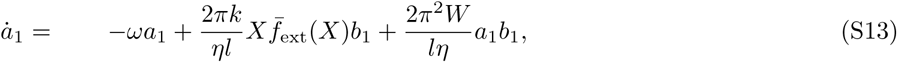

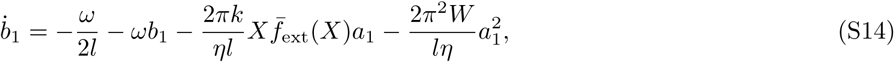

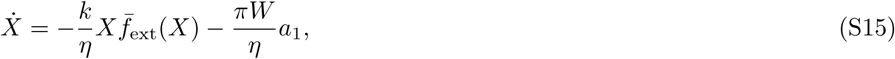

which, by adimensionalizing the time 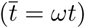 and length 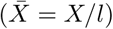, leads to:

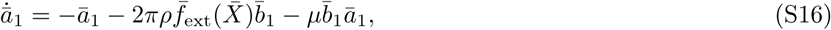

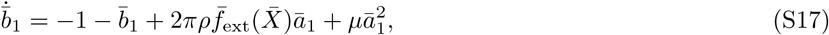

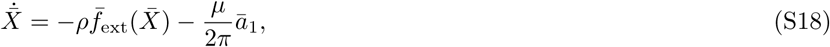

where 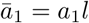 and *ρ* (resp. *µ*) quantify the ratio of an elastic recoil velocity *v_k_* = (*kl*)*/*(2*πη*) (resp. the motor velocity *v*_1_ = *πW/*(*lη*)) by the spontaneous velocity *v*_0_ = *ωl/*(2*π*):

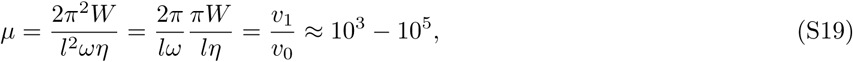

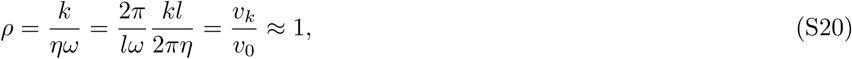

Numerical estimates are based on the values reported in Table III. Due to the low value of the transition rate *ω*, the the activity parameter *µ* is large. In the next paragraph, we show that this large of *µ* is responsible for the observation of *l/*2 steps.

#### Amplitude of steps - effective potential

In this paragraph, we show that the mean field description of Eq. (S5) contains the physics of the half-actin period (*l/*2) steps. For simplicity, we focus on the case with no external force *ρ* = 0. From Eq. (S16), we obtain

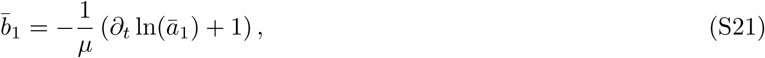

The substitution of the expression of 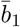 from Eq. (S21) into Eq. (S17) leads to

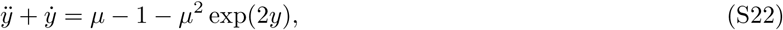

where 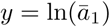 and 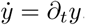. Eq. S22 corresponds to the dynamics of a particle in the following (effective) potential

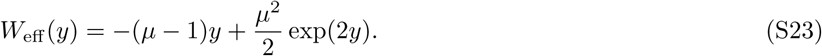

In the large *µ* limit, it takes typically *µ* oscillations for the damping to be effective. Thus in order to focus our attention on the periodic behavior characteristic of transients, we neglect the friction contribution *ẏ* in Eq. (S22). In this regime, we find that the following quantity (which is analogous to a total energy) is conserved:

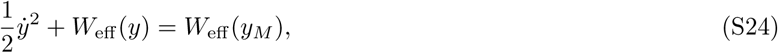

where *W*_eff_ (*y_M_*) is the energy at maximal value of *y* = *y_M_*; *W*_eff_ (*y_M_*) = *W*_eff_ (*y_m_*), where *W*_eff_ (*y_m_*) is energy at the minimal value of *y* = *y_m_* since the effective kinetic energy 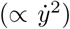 is zero both at *y_m_* and *y_M_* (by definition). The relation of Eq. (S24) leads to the following integrated dynamics on *y*

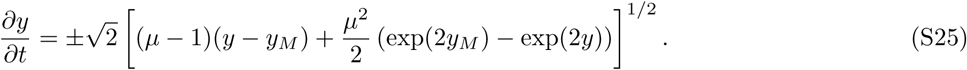

Integrating the force balance equation Eq. (S18), with *ρ* = 0, we find that the traveled distance 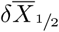 during a half period that spans from *y* = *y_m_* at time *t_m_* to *y_M_* at time *t_M_* reads:

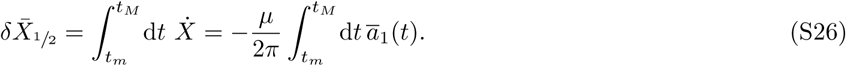

We then perform the change of variable from *t* to *y*, whose Jacobian is given by Eq. (S25), and then from 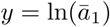 to 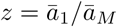. We obtain:

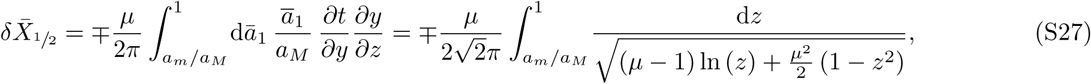

where the sign *∓* depends on the sign of the derivative 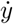. In the limit *µ → ∞*, the integral limit *a_m_/a_M_ →* 0. Based on the identity 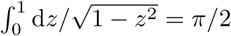, we find that Eq. (S27) converges to 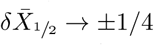, in unit of the actin period *l*. The total distance covered during a full step (denoted 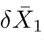) corresponds to twice this value: 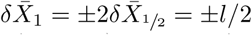. The sign ± correspond to the fact the steps are identical during the relaxation (negative velocity) of the contraction phase (positive velocity).

In conclusion, we have shown that, during transients, motors perform half-period steps in the limit of a large activity parameter *µ → ∞*, which corresponds to the limit of low transition rate *ω →* 0.

#### Force/velocity relation

We can estimate the mean force exerted by the assembly of motors for a given fixed velocity 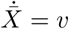 by determining the fixed point associated to Eqs. (S16–S17). This corresponds to the *adiabatic* approximation [11], which consists in that the modes *a*_1_ and *b*_1_ instantaneously relax to their steady state values. Under this assumption and the imposed constraint 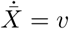, we obtain that the characteristic motor force behaves as

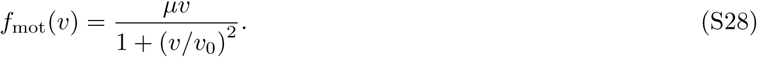

In the absence of an external force (*f*_ext_ = 0), the motor assembly reaches a steady state velocity 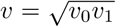, where *v*_0_ and *v*_1_ are defined in Eq. (S19) and Eq. (S20). Searching for extrema of the force-velocity relation Eq. (S28), we find that the maximal force that can be achieved by the collective motor assembly reads:

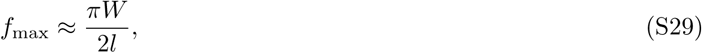

in the limit *v*_0_ ≫ *v*_1_. With *W* = 40 *k_b_T* (see Eq. (S3)) and *l* = 5 nm, we find that the maximal force per motor that can be exerted by the assembly is in the *f*_max_ ≈ 60 pN range. This expression is consistent with the estimate proposed in Eq. (2).

---

**Figure S1:**
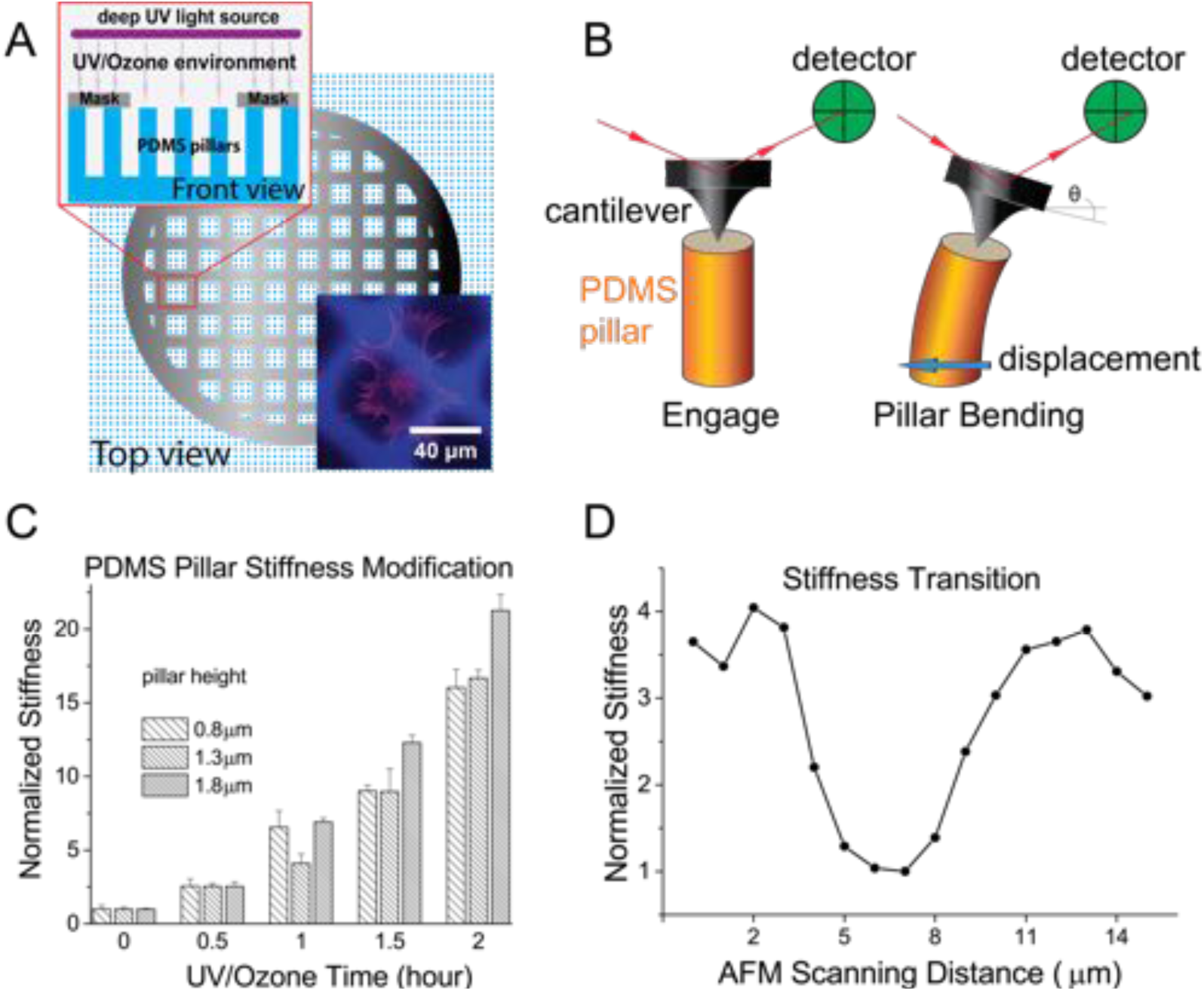
Dual Rigidity Pillar Platform Implementation and Characterization. A. Nickel TEM grid is placed on PDMS pillar substrate for UV/Ozone treatment (wavelengths: 185 nm and 254 nm). PDMS is premixed with c343 dye for rigidity marking. Inset: a MEF stained with phalloidin straddles multiple stiffness boundaries. Blue fluorescence signal is from c343 and marks the pillars that were in the mask shadow, and hence remain soft. B. Pillar bending stiffness increase is characterized by AFM in force volume and contact mode C. Stiffness increase was analyzed for 0.8, 1.3, and 1.8 *µ*m tall pillars. Stiffness increases exponentially with UV/Ozone treatment dose. Stiffness of the 1.8 *µ*m pillars increases up to 20 fold when treated for 2 hours. D. Stiffness changes finish the transition within 3 *µ*m distance.

**Figure S2:**
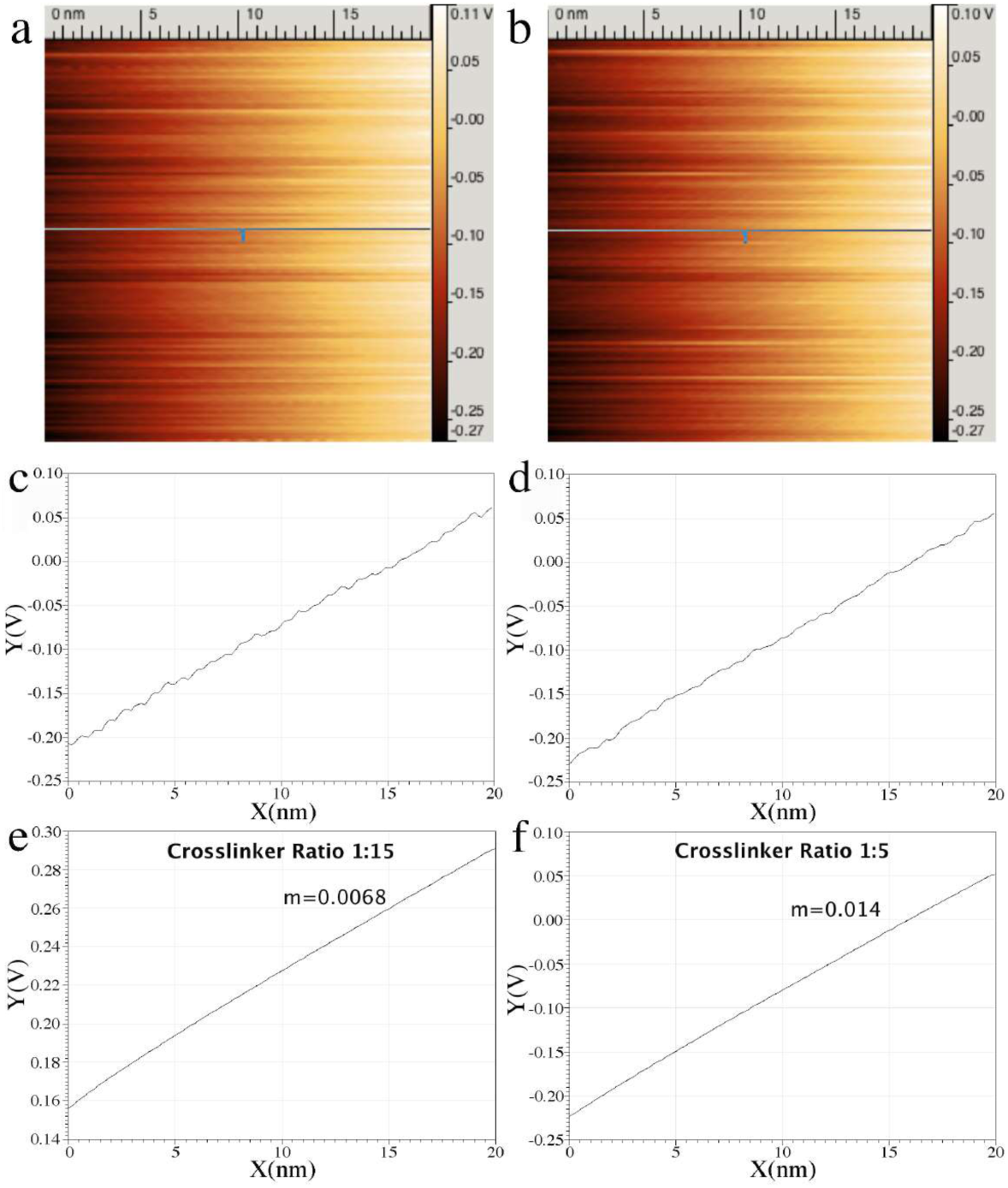
Direct AFM measurement of pillar bending stiffness. AFM friction channel output showing the voltage during the traces (bending, a) and retraces (unbending, b) of multiple scans of a single pillar. Each horizontal line represents a single scan/rescan, with the scanning distance marked by the scale bar at top. Voltage output is color coded with the range legend to the right of the image. Single scans from (a) and (b) (marked by grey line) are shown in (c) and (d), respectively. Linear voltage vs displacement relationship shows that pillars are acting as Hookian springs. Direct measurement of bending stiffness shows expected changes in spring constant. High crosslinker ratio PDMS (f) shows the 2-fold increase in bending stiffness compared to low crosslinker ratio PDMS (e) that is predicted due to the 2-fold change in bulk elastic modulus (note different y-axis scale).

**Figure S3:**
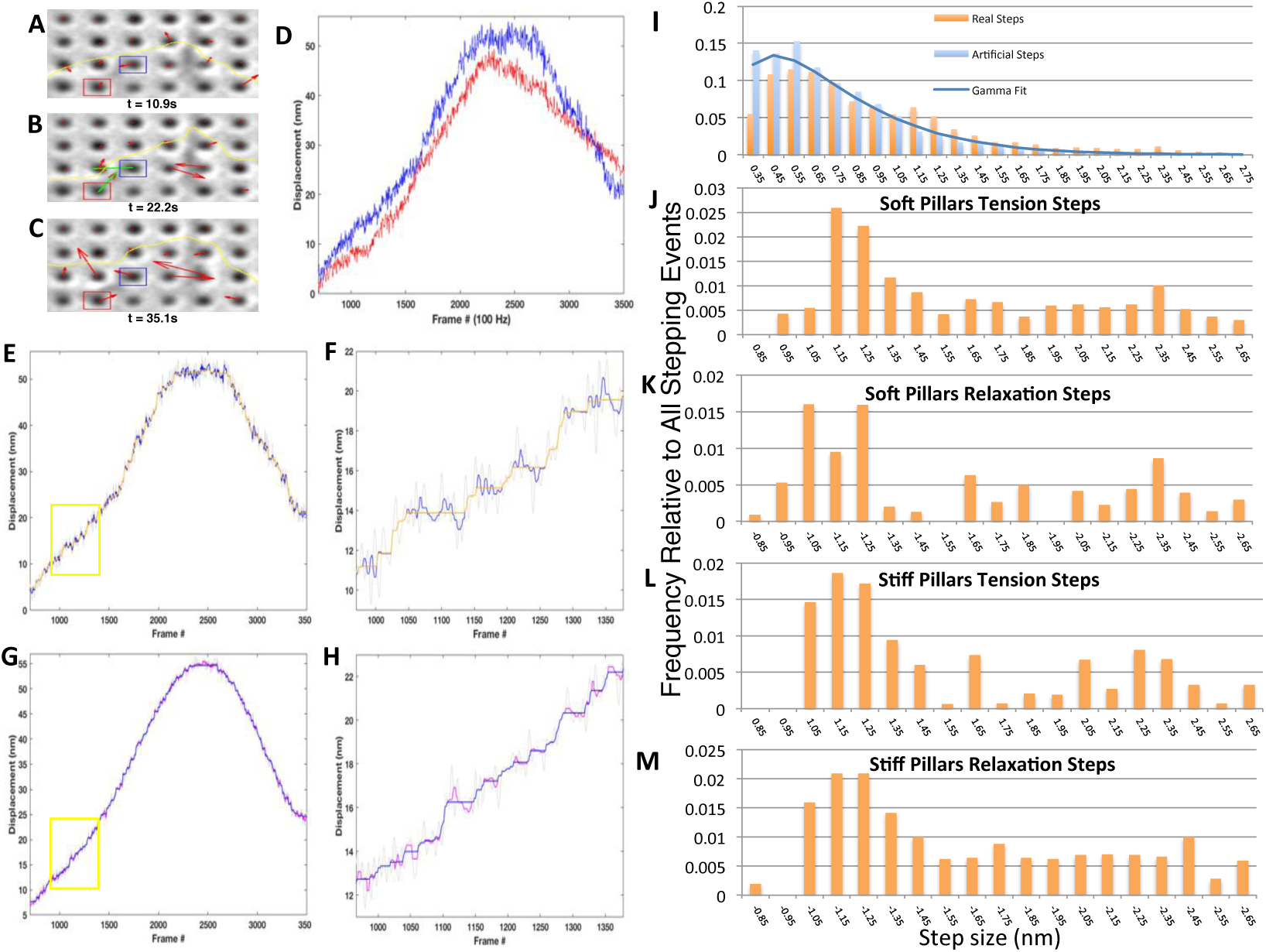
Characterization of Myosin Stepping Events. A,B,C. Snap shots from the beginning, peak, and end of a contractile unit. D. Displacement traces of the two pillars in A,B,C. Blue boxed pillar corresponds to the blue displacement curve; red boxed pillar corresponds with the red displacement curve. E. Blue displacement curve from D fitted with the l1-pwc curve fitting algorithm [44]. Grey represents raw displacement data; blue represents median filtered data; orange represents fitted steps. F. Zoom-in on yellow box from E. G. Artificial displacement curve generated by overlaying the noise data on top of a polynomial fit of the curve in E. Grey is raw data. Magenta is 15 point median filtered data. Blue line represents the output of running this generated data through the same step-fitting algorithm H. Zoom in on the yellow box in G. I. Histogram of steps generated by fitting displacements curves for contractile units on soft pillars. Real data in orange, artificial data in blue. J. Gamma distribution fit of blue data in I subtracted off of orange distribution in I revealing steps from our real data that cannot be due to noise (soft pillars, rising steps). K. Gamma fit of artificial distribution subtracted off of real distribution for falling steps on soft pillars. L. Gamma fit of artificial distribution subtracted off of real distribution for rising steps on stiff pillars. M. Gamma fit of artificial distribution subtracted off of real distribution for falling steps on stiff pillars. Notice: all distributions have a peak around 1.2 nm. For each condition *N >* 767 steps from *>* 11 cells across 5 different experiments.

**Figure S4:**
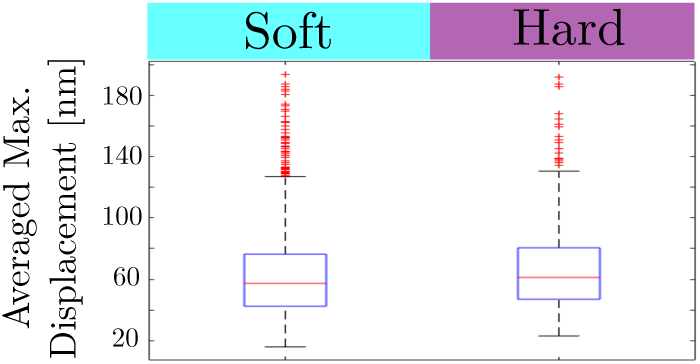
Average maximum displacements for contractile units formed over soft pillars (*K* = 3 pN.nm^*−*1^) and hard pillars (*K* = 60 pN.nm^*−*1^). We observe no statistical difference between the soft and hard populations (*n >* 500 for each condition from *>* 20 total cells over 5 different experiments).

**Figure S5:**
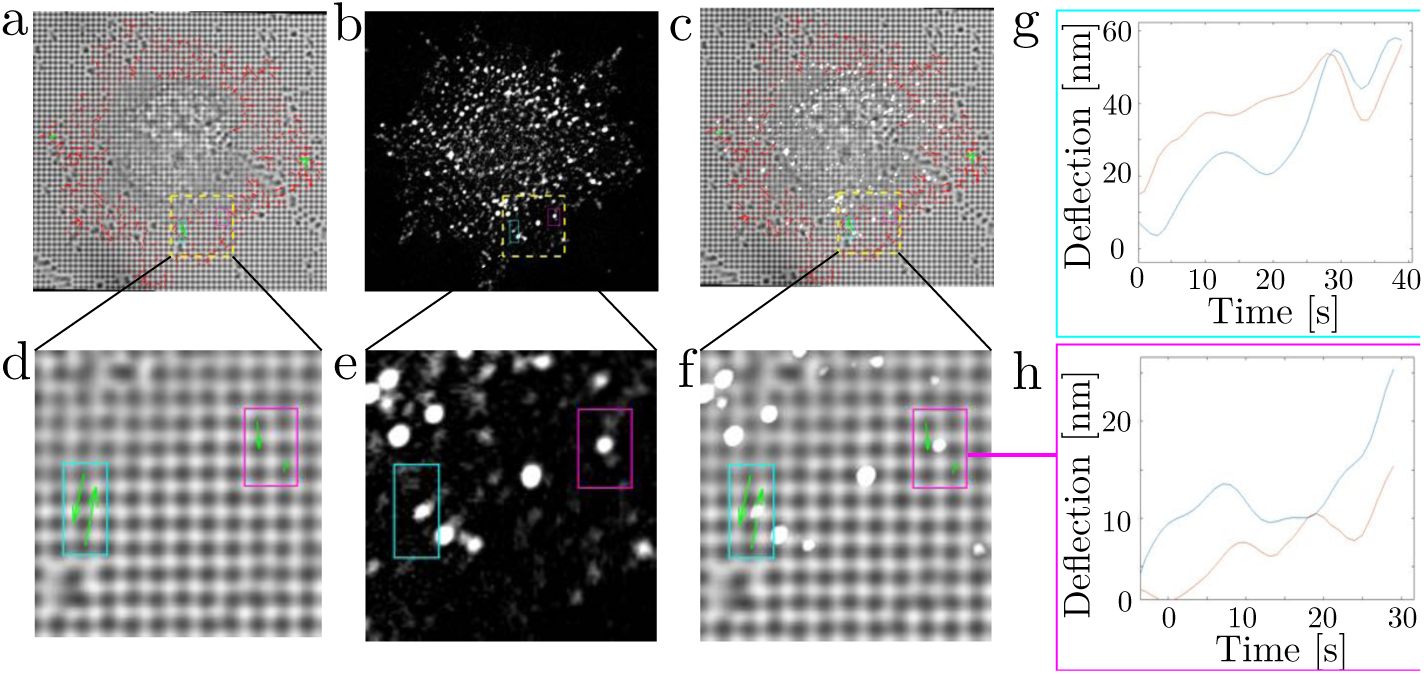
Contractile units on soft pillars. (a) Displacement vectors overlaid on a brightfield image of a spreading fibroblast immediately before fixation: (red arrows) pillar displacement vectors (green arrows) displacement vectors for anti-correlated pillars. (b) Fluorescent image of the same cell with staining of the biphosphorylated myosin regulatory light chain. (pMLC). (c) Merge of (a) and (b). (d-f) Blow up on the dashed yellow boxes in (a-c). (g-h) Pillar traces associated with the highlighted contractile units: (g) corresponds to the pillars within the cyan box in (f) and the associated myosin signal contains 44 activated heads. (h) tracks the pillars within the magenta box in (f) and contains 70 activated heads. Both have one myosin mini-filament responsible for force generation.

**Figure S6:**
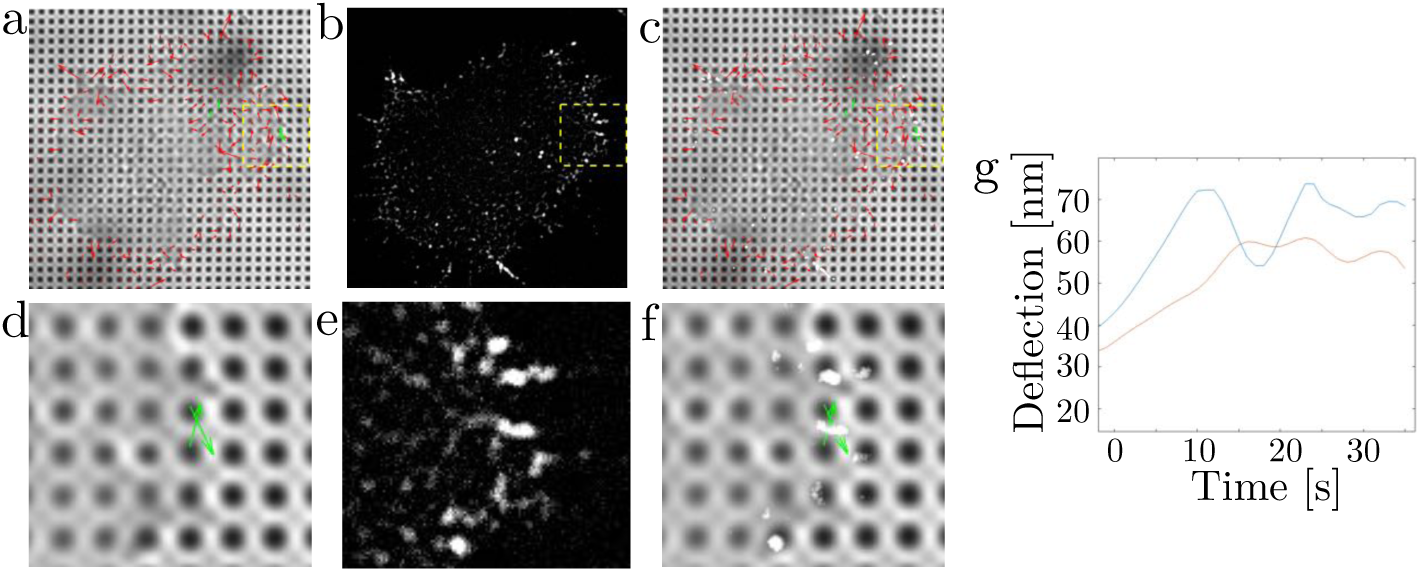
Contractile units on hard pillars. (a) Displacement vectors overlaid on a brightfield image of a spreading fibroblast immediately before fixation: (red arrows) pillar displacement vectors (green arrows) displacement vectors for anti-correlated pillars. (b) Fluorescent image of the same cell with staining of the biphosphorylated myosin regulatory light chain. (pMLC). (c) Merge of (a) and (b). (d-f) Blow up on the dashed yellow boxes in (a-c). (g-h) Pillar traces associated with the highlighted contractile units: (g) corresponds to the pillars within the cyan box in (f) and the associated myosin signal contains 44 activated heads. (h) tracks the pillars within the magenta box in (f) and contains 70 activated heads. Both have one myosin mini-filament responsible for force generation.

**Figure S7:**
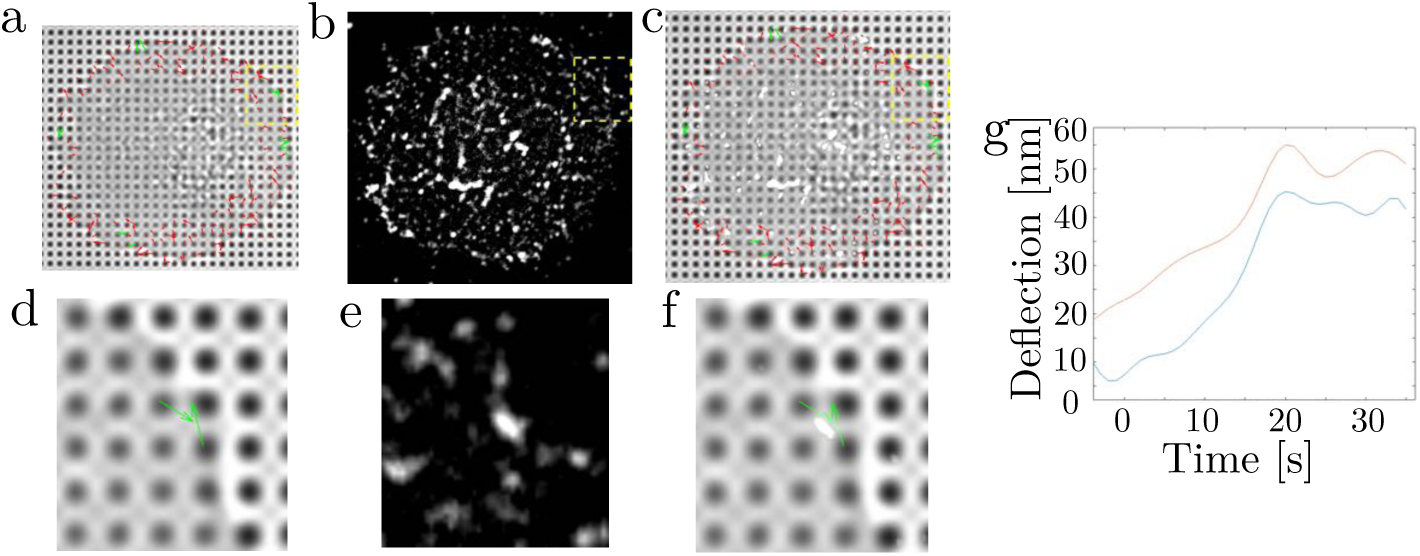
Contractile units on hard pillars. (a) Displacement vectors overlaid on a brightfield image of a spreading fibroblast immediately before fixation: (red arrows) pillar displacement vectors (green arrows) displacement vectors for anti-correlated pillars. (b) Fluorescent image of the same cell with staining of the biphosphorylated myosin regulatory light chain. (pMLC). (C) Merge of (a) and (b). (d-f) Blow up on the dashed yellow boxes in (a-c). (g-h) Pillar traces associated with the highlighted contractile units: (g) corresponds to the pillars within the cyan box in (f) and the associated myosin signal contains 44 activated heads. (h) tracks the pillars within the magenta box in (f) and contains 70 activated heads. Both have one myosin mini-filament responsible for force generation.

**Figure S8:**
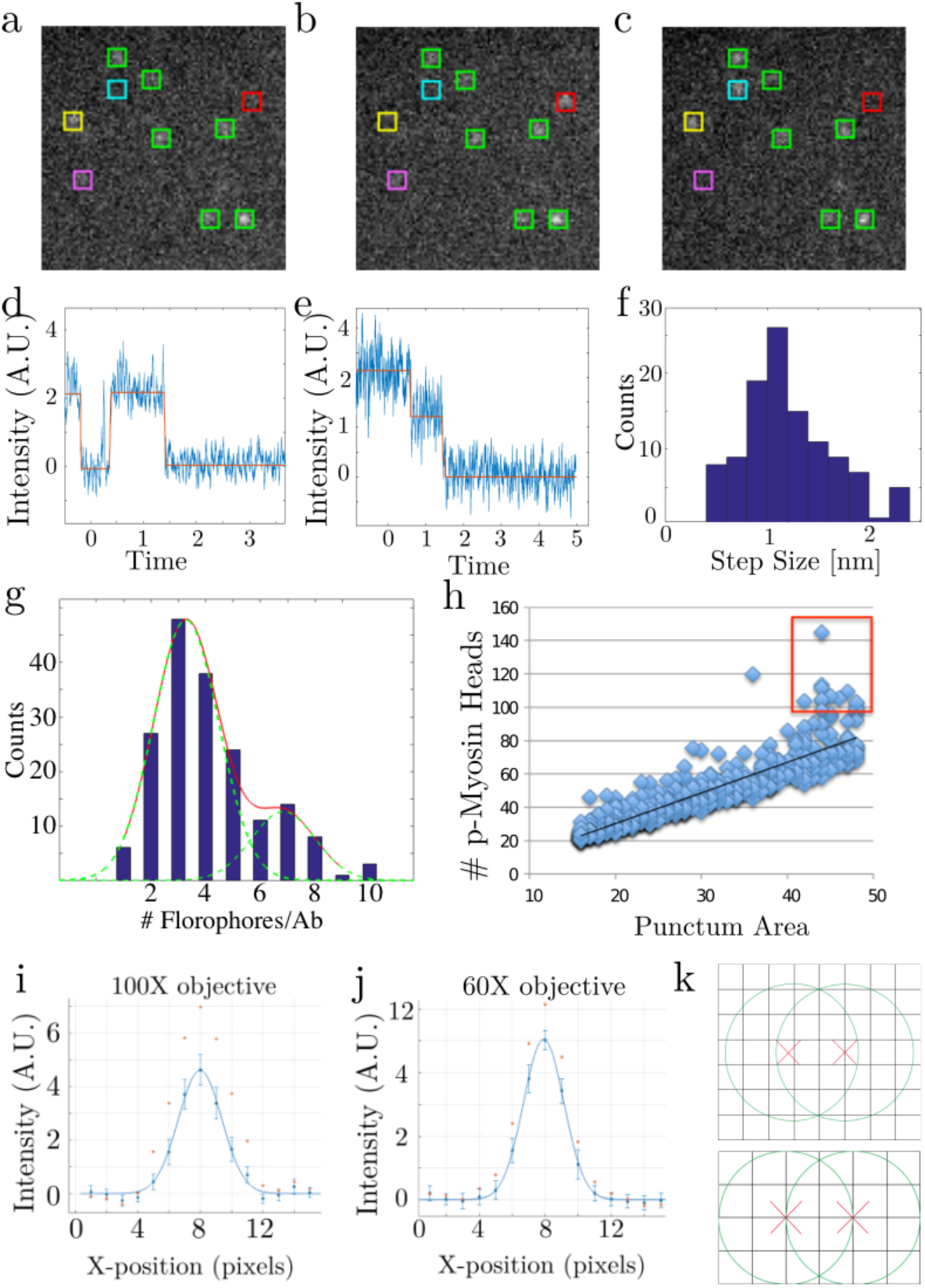
Fluorescence analysis. (a,b,c) Sequential images from a bleaching series of single secondary antibodies. Each image is the average of 50 consecutive 20 ms exposures. Green boxes show fluorophores that are emitting throughout. Yellow box shows a flurophore blinking off and then back on. Red box shows an fluorophore blinking on and then back off. Purple box shows a fluorophore bleaching out. (d) Intensity curve of a blinking fluorophore. The equivalence of the on and off steps show the uniform intensity of the fluorophore. Time unit: 1 s. (e) Sequential bleaching events from two fluorophores attached to the same antibody. (f) Histogram of bleaching events from the 100X objective shows the same prominent peak for single events and small peak for double events that we see from our 60X experiments, validating the technique. (g) Histogram of the number fluorophores conjugated to a single secondary antibody (denoted Ab), determined by dividing the signal from a single antibody prior to any bleaching by the fluorescence of a single alexa-647 fluorophore. Fitted gaussians peak at 3.3, 6.8, and 10 representing single antibodies and aggregations of two or three antibodies, respectively. (h) Relationship of punctum area (and hence filament length) with number of myosin heads is linear, except for some outliers at the upper end of the area range (red box) that are likely stacked filaments. (i-j) Analysis of the signal due to a point source emitter (single secondary antibodies) for the 100X and 60X objectives. Curve generated by taking a line scan over 40 different point sources, averaging the line scans together, and fitting a Gaussian in matlab. Red stars mark the curve from the raw images, blue circles mark the curve from rolling ball filtered images. These point spreads were used to calculate the puncta intensity from the particle analysis output from ImageJ. (k) Representations of the largest and small possible area for a single filament (respectively) based on the maximum single pixel intensity observed in filaments, the point spread function of the 60X objective, and the spacing of poles in a bipolar filament. These areas were used to bound the puncta that were included in the measurement of the number of molecules in a single filament. Puncta smaller than 24 pixels are unlikely to be bipolar filaments (They are perhaps the remnants of disassembled filaments). Puncta larger than 48 pixels are taken to be assemblies of multiple filaments (and are more prevalent over more rigid substrates).

**Figure S9:**
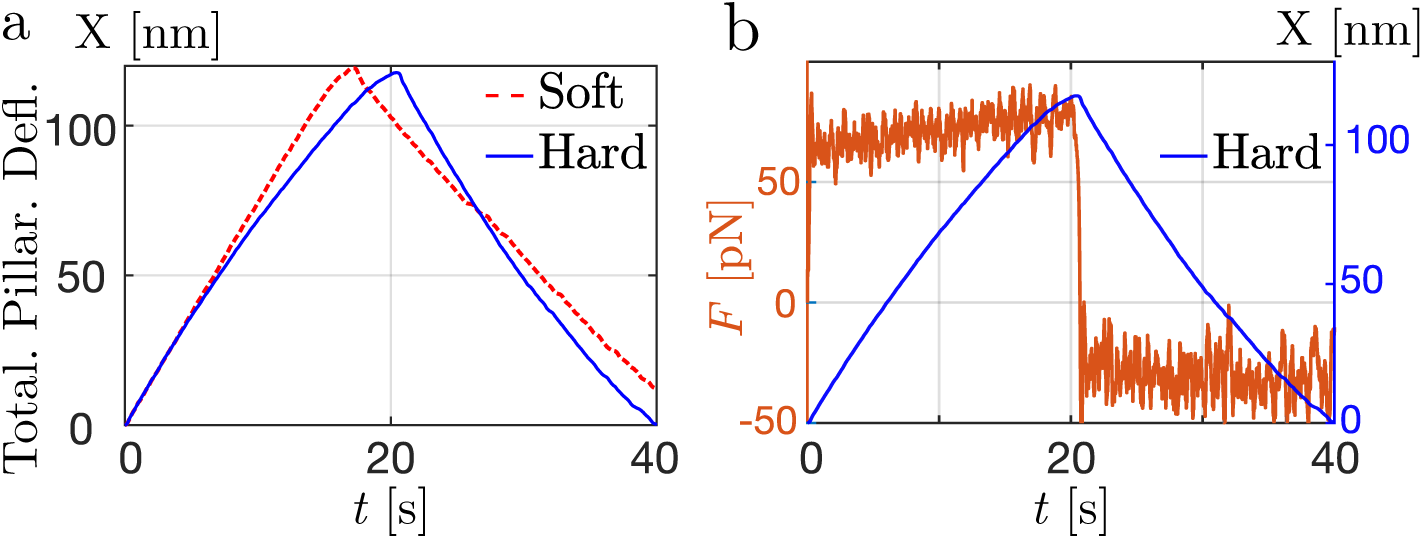
(a) Simulations predicting the total pillar deflection *X*(*t*) for hard pillars (solid blue curve; *K* = 60 pN.nm^*−*1^, *N* = 160) and soft pillars (dashed red line; *K* = 3 pN.nm^*−*1^, *N* = 120). Other parameters are specified in Tab. III. (b) Evolution of the force generated per motor as a function of time (orange curve), superposed with the evolution of the pillar deflection (dark blue curve) for hard pillars. The short-time oscillations in the force correspond to steps in the pillar deflection.

**Figure Movie 1:**
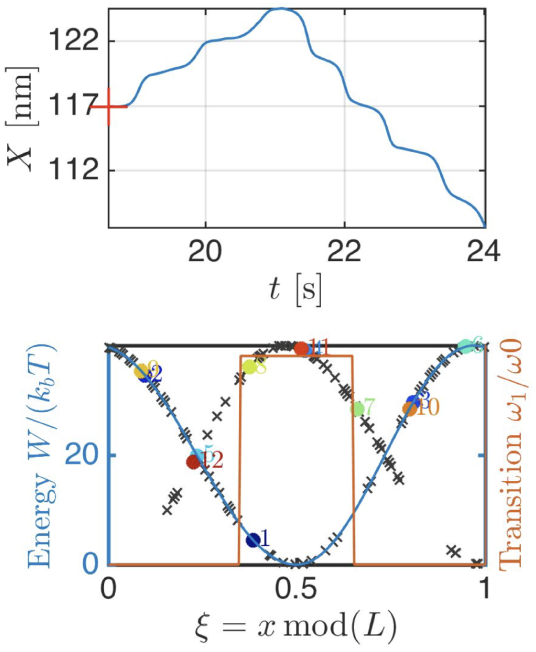
Stochastic simulation for a transition rate localized around the minimum of the potential: (top panel) total pillar deflection as a function of time. The red cross indicate the current state of the assembly. (bottom panel) State of the motors assembly (color symbols) cyclic position *ξ_i_*= mod (*x_i_, l*) and energy of a particular motor (blue curve) potential *W*_1_ in state 1 as a function of the cyclic coordinate *ξ* (orange curve) transition rate *ω*_1_ from state 1 to state 2. Parameters are *K* = 10 pN.nm^*−*1^ and *N* = 120 (other parameters are given in Tab. III).

**Figure Movie 2:**
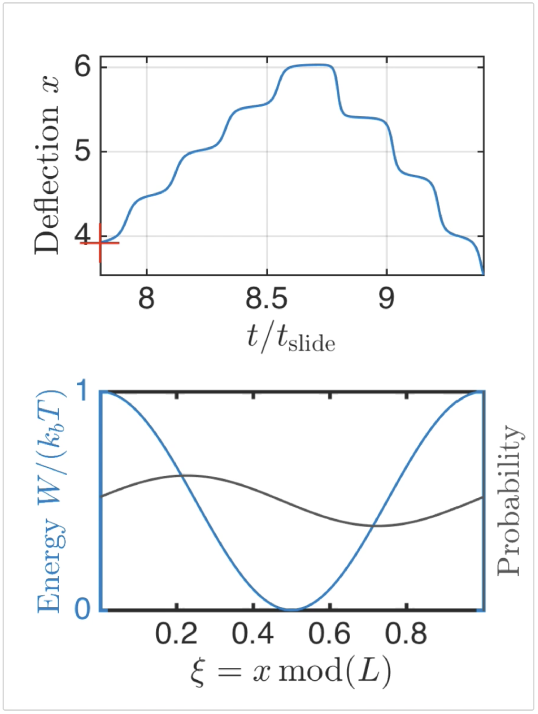
Analytical resolution for a constant total transition rates *ω* of main text Eq. (3) corresponding to the large number of motor limit: (top panel) total pillar deflection *X*(*t*) of the collective assembly as a function of time *ωt*. The red cross indicate the current value of the pillar deflection. (bottom panel) Joint energy-density representation – (blue curve) energy ^profile in state 1 as a function of the cyclic coordinate *ξ* = mod (*x, l*) (grey curve) occupation density *P*_1_ in state 1. The occupation density in state 2 reads *P*_2_= 1*/l − P*_1_.^

**Table I:** UV/Ozone adjusted PDMS rigidity as measured by AFM (bending stiffness) and calculated from bending stiffness (effective stiffness).

| PDMS Pillar Height (μm) | Bending Stiffness (pN/NM) | Bending Stiffness (2 hr UV/Ozone) | Effective Stiffness (kPa) | Effective Stiffness (2 hr UV/Ozone) |
| --- | --- | --- | --- | --- |
| 1.8 | 3 | 60 | 9 | 170 |
| 1.3 | 6 | 100 | 18 | 310 |
| 0.8 | 19 | 310 | 55 | 910 |

**Table 2:**
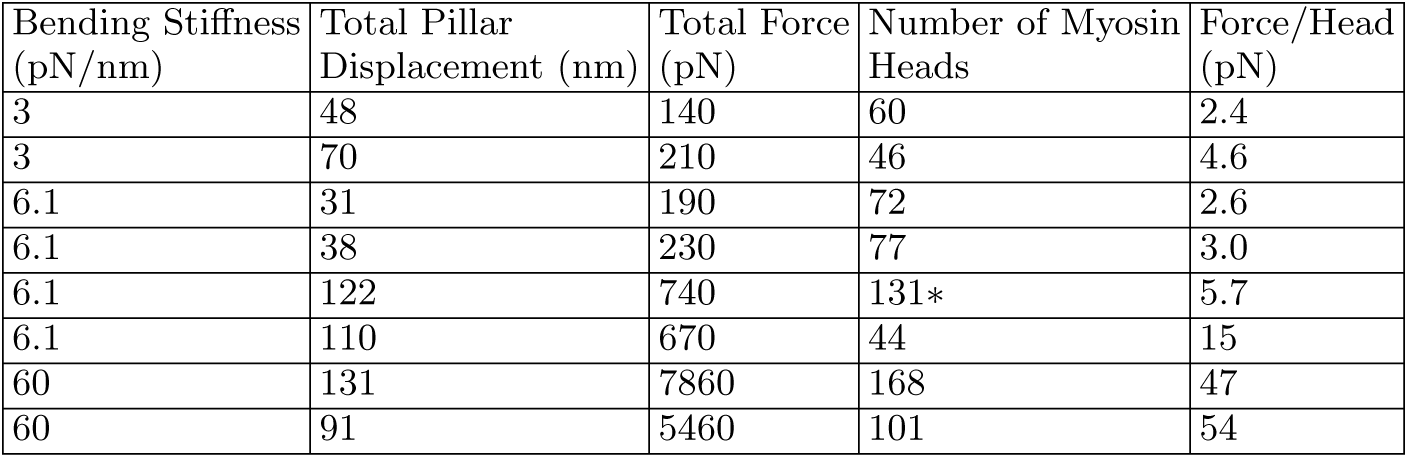
Myosin Filaments Associated with Active Sarcomeric Units. Mulitple myosin filaments were associated with peak displacements on stiff pillars, and with contractile units made by more than two pillars on softer pillars. Notably, all captured contractions that could be associated with p-Myosin filaments were either in the contraction phase, or at the peak contraction displacement, suggesting myosin phosphatase may trigger the relaxation phase of the contractile unit. - signifies a contractile unit formed by three pillars. Variable rigidities are obtained using the following procedure: 3 pN.nm^*−*1^: 1.8 *µ*m pillars, no UV; 6.1 pN.nm^*−*1^: 1.3 *µ*m pillars, no UV; 60 pN.nm^*−*1^: 1.8 *µ*m pillars, 2 hour UV treatment.

| Bending Stiffness<br>(pN/nm) | Total Pillar<br>Displacement (nm) | Total Force<br>(pN) | Number of Myosin<br>Heads | Force/Head<br>(pN) |
| --- | --- | --- | --- | --- |
| 3 | 48 | 140 | 60 | 2.4 |
| 3 | 70 | 210 | 46 | 4.6 |
| 6.1 | 31 | 190 | 72 | 2.6 |
| 6.1 | 38 | 230 | 77 | 3.0 |
| 6.1 | 122 | 740 | 131* | 5.7 |
| 6.1 | 110 | 670 | 44 | 15 |
| 60 | 131 | 7860 | 168 | 47 |
| 60 | 91 | 5460 | 101 | 54 |

**Table 3:** List of notations and parameters estimates.

| Quantity | Meaning | Value |
| --- | --- | --- |
| $l$ | Actin periodicity | 5 nm [29] |
| $\omega_0$ | Transition rate | 5 Hz |
| $\eta$ | Friction (per motor) | $2 \cdot 10^{-3} \text{ N.s.m}^{-1}$ |
| $W$ | Potential energy | $40 k_b T$ |
| $v_1 = \pi W/(l\eta)$ | Sliding velocity | $10 \mu\text{m.s}^{-1}$ |
| $K$ | Pillar elasticity | $3 - 60 \text{ pN.nm}^{-1}$ |
| $N$ | Number of heads | 110 – 170 |
| $e$ | Width of the transition zone | 1 nm |
| $\delta$ | Asymmetry of the potential | 0.30 nm |

